# Uniquely excitable neurons enable precise and persistent information transmission through the retrosplenial cortex

**DOI:** 10.1101/673954

**Authors:** Ellen K.W. Brennan, Shyam Kumar Sudhakar, Izabela Jedrasiak-Cape, Omar J. Ahmed

**Author notes:** Co-first authors (Ellen K.W. Brennan, Shyam Kumar Sudhakar). **CORRESPONDENCE TO:** Omar J. Ahmed, *Mail:* Dept. of Psychology, 530 Church St. University of Michigan, Ann Arbor, MI 48109, *Email:.

## Abstract

The retrosplenial cortex (RSC) is essential for both memory and navigation, but the neural codes underlying these functions remain largely unknown. Here, we show that the most prominent cell type in layers 2/3 (L2/3) of the granular RSC is a uniquely excitable, small pyramidal cell. These cells have a low rheobase (LR), high input resistance, lack of spike-frequency adaptation, and spike widths intermediate to those of neighboring fast-spiking (FS) inhibitory neurons and regular-spiking (RS) excitatory neurons. LR cells are excitatory but rarely synapse onto neighboring neurons. Instead, L2/3 of RSC is an inhibition-dominated network with dense connectivity between FS cells and from FS to LR neurons. Biophysical models of LR but not RS cells precisely and continuously encode sustained input from afferent postsubicular head-direction cells. Thus, the unique intrinsic properties of LR neurons can support both the precision and persistence necessary to encode information over multiple timescales in the RSC.

## INTRODUCTION

The retrosplenial cortex (RSC) plays a critical role in learning and memory. In humans, damage to the RSC via hemorrhage or tumor results in both anterograde and retrograde amnesia, often purging several years of recent memories (Ironside and Guttmacher, 1929; Heilman and Sypert, 1977; Valenstein et al., 1987; Todd and Bucci, 2015; Chrastil, 2018). Similar impacts on both anterograde and retrograde memory are also seen in monkeys when the RSC is lesioned (Buckley and Mitchell, 2016). In rodents, RSC lesions impair performance on both spatial learning and fear conditioning tasks (Vann et al., 2003, 2009; van Groen et al., 2004; Keene and Bucci, 2008; Katche et al., 2013; Todd et al., 2015, 2017; Sigwald et al., 2016; Yamawaki et al., 2019b). Recent imaging studies in mice confirm that RSC neurons can display evidence of long-duration, persistent spatial memory engrams (Czajkowski et al., 2014; Milczarek et al., 2018; de Sousa et al., 2019; Hattori et al., 2019).

The RSC is also critical for spatial navigation (Maguire, 2001; Epstein, 2008). Human case studies show that damage to the RSC leads to disorientation in space in addition to memory impairments (Bottini et al., 1990; Takahashi et al., 1997; Ino et al., 2007; Osawa et al., 2007). Such patients can identify known scenes or locations but are unable to extract any orientation or location information from them and thus experience difficulties navigating even familiar environments (Bottini et al., 1990; Takahashi et al., 1997; Ino et al., 2007). A neuroimaging study identified the coding of head direction information in the RSC while participants navigated a novel virtual environment, suggesting that the visual cues of orientation are processed in part by the RSC during navigation (Shine et al., 2016). Animal studies also report encoding of spatial information within the RSC, including that of head direction, position, and turning behavior (Cho and Sharp, 2001; Alexander and Nitz, 2015; Vedder et al., 2016; Mao et al., 2017, 2018).

How is the RSC uniquely suited to carry out these spatial memory and navigation computations? This is a fundamental but unsolved circuit input-output transformation problem. The RSC receives prominent spatial and memory-related inputs from the hippocampus, subicular complex, anterior thalamus, secondary motor cortex, and visual cortex, as well as the contralateral RSC (van Groen and Wyss, 1990, 2003; Wyss and van Groen, 1992; Miyashita and Rockland, 2007). Recent studies have started to document the functional nature of these inputs to the RSC (Yamawaki et al., 2016, 2019a, 2019b; Sempere-Ferràndez et al., 2018; Sempere-ferràndez et al., 2019). However, the precise properties of the RSC neuronal subtypes involved (Wyss et al., 1990; Sugar et al., 2011; Kurotani et al., 2013) is rarely studied and the local connectivity between RSC subtypes completely unknown. While attractor network models of RSC incorporating generic neurons exist (Bicanski and Burgess, 2016; Page and Jeffery, 2018), it is critical to discover the key local intrinsic and synaptic properties that allow RSC to perform its specialized functions. Without this information, it is impossible to develop biophysically realistic models of RSC cells or circuits, which would in turn help to decipher the exact coding schemes being employed by the RSC.

Here, we investigate the intrinsic physiology, local synaptic connectivity, and computational abilities of cells within the superficial layers of granular retrosplenial cortex (RSG). The majority of neurons within this region are a distinct subtype of small, highly excitable, non-adapting pyramidal neurons. We show, for the first time, that these cells are excitatory but, surprisingly, rarely excite their neighboring inhibitory or excitatory neurons. Instead, there is prevalent local inhibition from fast-spiking (FS) L2/3 neurons onto these highly excitable neurons and between pairs of FS cells, highlighting a network dominated by feedforward, not feedback, inhibition. We then use this information to construct biophysically-realistic computational models of RSC cell types and investigate how they process realistic, *in vivo* spike trains of incoming information. We find that the uniquely excitable principal neurons in the RSG are optimally suited to precisely and persistently encode the sustained head-direction input they receive from the postsubiculum. A smaller population of regular-spiking (RS) excitatory neurons in L2/3 show pronounced adaptation and are unable to maintain such sustained, high-frequency responses. Our results show that there are two complementary coding strategies operating in parallel in the superficial retrosplenial cortex.

## RESULTS

### Low Rheobase cells are highly excitable neurons in the superficial retrosplenial granular cortex

We recorded from 167 cells in the superficial layers (L2/3) of the mouse retrosplenial granular cortex (RSG). Consistent with other cortical regions, fast-spiking (FS) interneurons were present in these RSG layers (Figure 1A). FS cells were identified by their unique spiking properties (Connors and Gutnick, 1990; Sempere-Ferràndez et al., 2018), including their narrow spike width and rapid, sharp AHPs. Regular-spiking (RS) pyramidal neurons were occasionally found, but far less often than in typical neocortex (Figure 1B). A third population of cells was identified. The action potentials of these neurons were narrower than typical RS cells and often displayed prominent afterdepolarizations (Figure 1C). Detailed analyses of physiological and intrinsic parameters revealed several distinctly unique properties of these neurons. Statistical differences were calculated using a two-tailed Wilcoxon rank sum for all intrinsic comparisons. Spike widths were between those of FS and RS cells (FS = 0.22 ± 0.05 ms, RS = 0.86 ± 0.05 ms, LR = 0.55 ± 0.02ms; p<0.01 for each comparison; Figure 1D; Table 1). Additionally, these cells had uniquely high input resistance (402.69 ± 16.75 MΩ p<0.01), low input capacitance (38.42 ± 1.32 pF; p<0.01), and low rheobase (91.79 ± 12.89 pA; p<0.01), suggesting they are a class of highly excitable neurons distinct from both FS and RS neurons (Figure 1E-G; Table 1). They also exhibited minimal spike frequency adaptation (ratio of 1.26 ± 0.06), far lower than the substantial spike frequency adaptation shown by RS cells (ratio of 3.42 ± 0.58; p<0.01), highlighting their potential ability to fire trains of action potentials at high frequencies with minimal adaptation (Figure 1J; Table 1). For reasons investigated and explained in detail below, we refer to these unique neurons as Low Rheobase (LR) cells in the rest of this manuscript.

**Figure 1.**
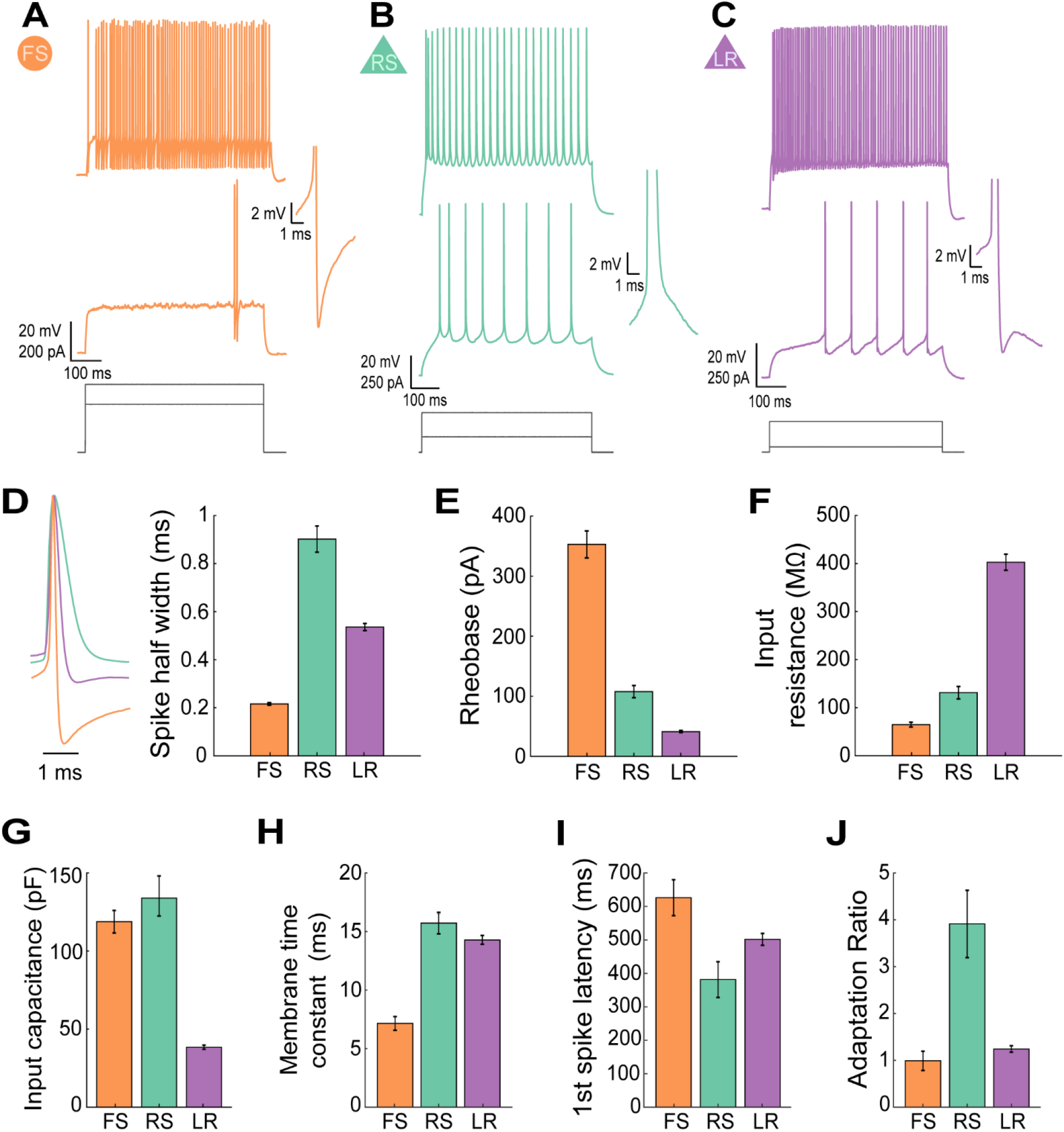
Low rheobase cells represent a highly excitable cell type in the superficial retrosplenial cortex. ***A.*** Intrinsic physiological properties of an FS neuron in the superficial layers of the granular retrosplenial cortex. Top trace, Ability to fire sustained high frequency trains of action potentials with little or no spike frequency adaptation. Middle trace, A substantial delay to first spike after current onset during a near-threshold current step. This late-spiking feature was consistent across all 42 recordings of FS cells in these layers. Right inset is a zoomed-in view of the first spike in the middle trace. It shows a narrow spike width followed by a large afterhyperpolarization. These features are distinctive of FS cells and were also consistent across all 42 recordings of FS cells in these layers. Bottom, injected current amplitudes for the voltage responses shown above. ***B.*** Intrinsic physiological properties of an RS neuron in the superficial layers of the granular retrosplenial cortex. Top trace, Presence of spike frequency adaptation when firing at higher frequencies in response to large suprathreshold current steps. This feature was consistent across all 17 recordings of RS cells in these layers. Middle trace, Minor delay to first spike after current onset during an at-threshold current step. Right inset is a zoomed-in view of the first spike in the middle trace. It shows a relatively wide spike width followed by a gradual return to baseline membrane potential. Bottom, injected current amplitudes for the voltage responses shown above. ***C.*** Intrinsic physiological properties of an LR neuron in the superficial layers of the granular retrosplenial cortex. Top trace, Ability to fire sustained high frequency trains of action potentials with little spike frequency adaptation. Middle trace, Moderate delay to first spike after current onset during an at-threshold current step. These features were consistent across the 108 recordings of LR cells in these layers. Right inset is a zoomed-in view of the first spike in the middle trace. It shows a moderate spike width followed by a clear afterdepolarization before returning to baseline membrane potential. This afterdepolarization was present in 66% of LRs recorded in these layers (n=61/93). Bottom, injected current amplitudes for the voltage responses shown above. ***D.*** Left, Representative traces from FS, RS, and LR cell action potentials overlaid to show differences in spike width. Right, Average spike widths showing that LR spike widths are intermediate to those of neighboring FS and RS [p<0.01 for each comparison; Wilcoxon rank sum]. ***E.*** Average rheobase for FS, RS, and LR cells showing a markedly low rheobase for LR cells compared to that of FS and RS [p<0.01 for each comparison; Wilcoxon rank sum]. ***F.*** Average input resistance (IR) for FS, RS, and LR cells showing a uniquely high IR for LR cells [p<0.01 for each comparison; two-tailed Wilcoxon rank sum]. ***G.*** Average input capacitance (IC) for FS, RS, and LR cells showing a markedly low IC for LR cells compared to FS and RS [p<0.01 for each comparison; Wilcoxon rank sum]. ***H.*** Average membrane time constant (tau) for FS, RS, and LR cells [LR v FS, p<0.001; LR v RS, p=0.1570; Wilcoxon rank sum]. ***I.*** Bar graph of the average latency to first spike after onset of an at-threshold current injection for FS, RS, and LR cells showing a substantial latency to first spike for both FS and LR cells in these layers [LR v FS, p=0.0089; LR v RS, p=0.0366; Wilcoxon rank sum]. ***J.*** Bar graph showing the average spike frequency adaptation ratio for FS, RS, and LR cells showing lack of adaptation in FS and LR cells [p<0.05 for each comparison; Wilcoxon rank sum]. Error bars represent standard error of the mean.

**Table 1.**
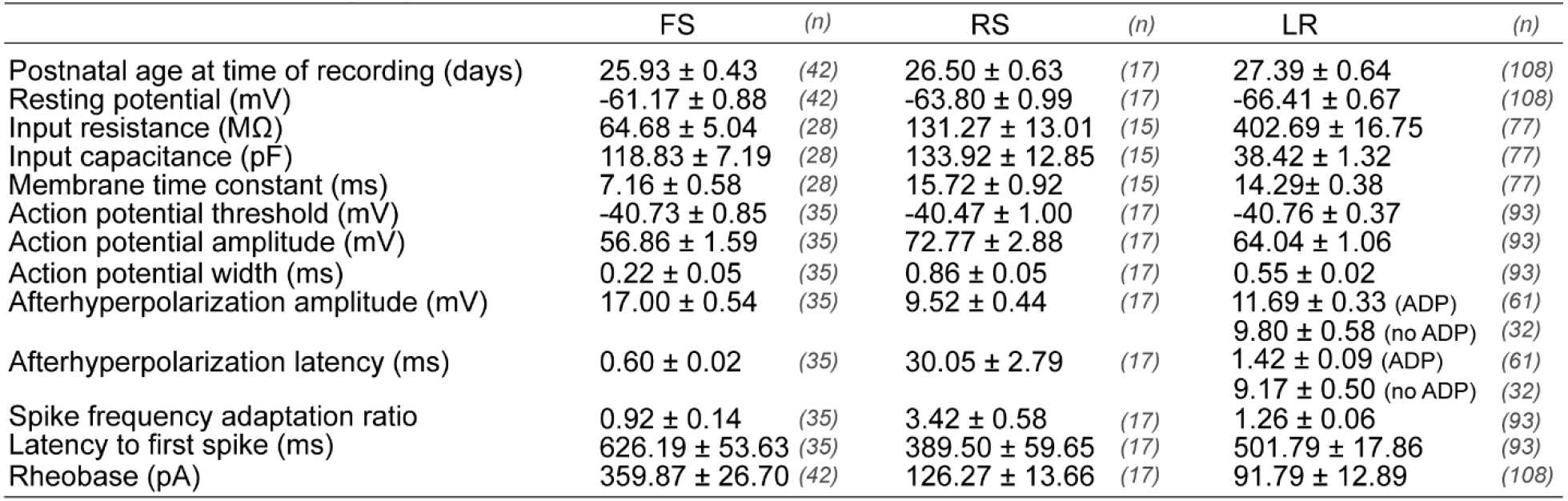
Intrinsic cell properties reveal that LR cells have a uniquely low rheobase, high input resistance, low input capacitance, and low spike frequency adaptation, as well as a spike width between that of FS and RS cells. Values are means ± SEM for each of the calculated intrinsic cell properties separated by cell group. Ns are reported individually for each property for each cell type. Details of the measurements are described in Methods. LR cells significantly differed from RS cells on the following measures: input resistance, input capacitance, spike width, spike frequency adaptation ratio, and rheobase (p<0.01) as well as latency to first spike (p<0.05). LR cells significantly differed from FS cells on the following measures: input resistance, input capacitance, membrane time constant, spike width, first spike latency, AHP amplitude, and rheobase (p<0.01), as well as spike frequency adaptation ratio (p<0.05). FS: Fast-spiking; RS: Regular-Spiking; LR: Low Rheobase.

|  | FS | (n) | RS | (n) | LR | (n) |
| --- | --- | --- | --- | --- | --- | --- |
| Postnatal age at time of recording (days) | $25.93 \pm 0.43$ | (42) | $26.50 \pm 0.63$ | (17) | $27.39 \pm 0.64$ | (108) |
| Resting potential (mV) | $-61.17 \pm 0.88$ | (42) | $-63.80 \pm 0.99$ | (17) | $-66.41 \pm 0.67$ | (108) |
| Input resistance (MΩ) | $64.68 \pm 5.04$ | (28) | $131.27 \pm 13.01$ | (15) | $402.69 \pm 16.75$ | (77) |
| Input capacitance (pF) | $118.83 \pm 7.19$ | (28) | $133.92 \pm 12.85$ | (15) | $38.42 \pm 1.32$ | (77) |
| Membrane time constant (ms) | $7.16 \pm 0.58$ | (28) | $15.72 \pm 0.92$ | (15) | $14.29 \pm 0.38$ | (77) |
| Action potential threshold (mV) | $-40.73 \pm 0.85$ | (35) | $-40.47 \pm 1.00$ | (17) | $-40.76 \pm 0.37$ | (93) |
| Action potential amplitude (mV) | $56.86 \pm 1.59$ | (35) | $72.77 \pm 2.88$ | (17) | $64.04 \pm 1.06$ | (93) |
| Action potential width (ms) | $0.22 \pm 0.05$ | (35) | $0.86 \pm 0.05$ | (17) | $0.55 \pm 0.02$ | (93) |
| Afterhyperpolarization amplitude (mV) | $17.00 \pm 0.54$ | (35) | $9.52 \pm 0.44$ | (17) | $11.69 \pm 0.33$ (ADP) | (61) |
| | | | | | $9.80 \pm 0.58$ (no ADP) | (32) |
| Afterhyperpolarization latency (ms) | $0.60 \pm 0.02$ | (35) | $30.05 \pm 2.79$ | (17) | $1.42 \pm 0.09$ (ADP) | (61) |
| | | | | | $9.17 \pm 0.50$ (no ADP) | (32) |
| Spike frequency adaptation ratio | $0.92 \pm 0.14$ | (35) | $3.42 \pm 0.58$ | (17) | $1.26 \pm 0.06$ | (93) |
| Latency to first spike (ms) | $626.19 \pm 53.63$ | (35) | $389.50 \pm 59.65$ | (17) | $501.79 \pm 17.86$ | (93) |
| Rheobase (pA) | $359.87 \pm 26.70$ | (42) | $126.27 \pm 13.66$ | (17) | $91.79 \pm 12.89$ | (108) |

In order to determine whether the LR neurons are a truly distinct neuronal subtype, we sought to identify the physiological properties that can clearly distinguish them from other neurons in the superficial RSG. Specifically, using features such as rheobase, input capacitance, input resistance, and spike width, we were able to isolate the LR cells from both FS and RS cells (Figure 2). LR cells cluster clearly and separately from FS and RS cells based on these intrinsic properties, suggesting they are a unique and distinct neuronal subtype compared to the others included in this study.

**Figure 2.**
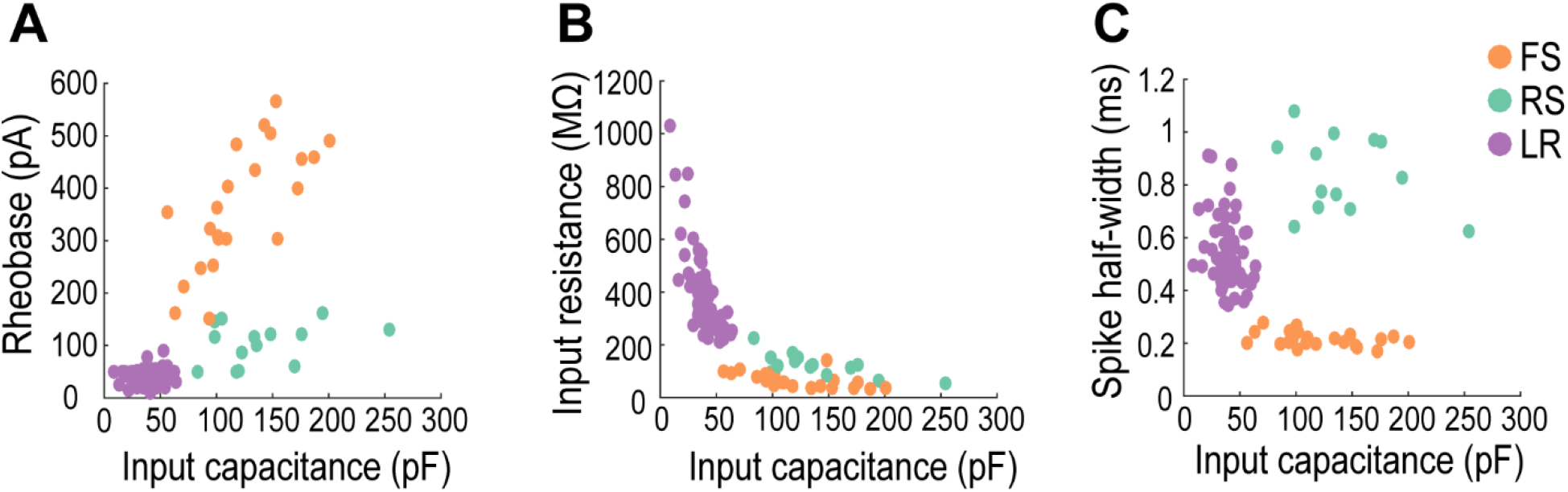
Low Rheobase neurons are a distinct, unique cell type. LR cells cluster clearly and separately from FS and RS cells when input capacitance is plotted against ***A.*** rheobase, ***B.*** Input resistance, and ***C.*** spike half-width.

### Low Rheobase cells are the dominant cell type in superficial retrosplenial granular cortex

LR neurons were the most commonly encountered cell type in L2/3 of RSG. To quantify the relative percentage of neurons in the superficial RSG, the recorded neurons were assigned to one of four groups based on their intrinsic physiological properties: FS, RS, LR, and unclassified. The unclassified group consisted of neurons whose intrinsic and/or firing properties did not fall under any of the three defined groups. Thus, this group includes other neuronal subtypes not investigated in this study as well as a few potential FS, RS, and LR neurons whose properties were difficult to classify. We found that LR cells are the dominant cell type in both layers 2 and 3, accounting for 51.9% of the neurons in layer 2 and 57.14% in layer 3 (Figure 3). However, 0 out of 25 recordings in layers 5 and 6 were of LR cells, suggesting that LR neurons are restricted to the superficial layers of RSG (data not shown). Surprisingly, the prevalence of RS cells in L2/3 of RSG was strikingly low, representing only 18.5% of all layer 2 neurons and 8.9% of layer 3 neurons. The FS probabilities are slightly skewed, as experiments detailed later in this manuscript specifically targeted FS interneurons. Thus, their probability reported here is likely slightly larger than their true representation in the cortex. Nonetheless, it is clear that LR cells are the most prevalent cell type within the superficial layers of the granular retrosplenial cortex.

**Figure 3.**
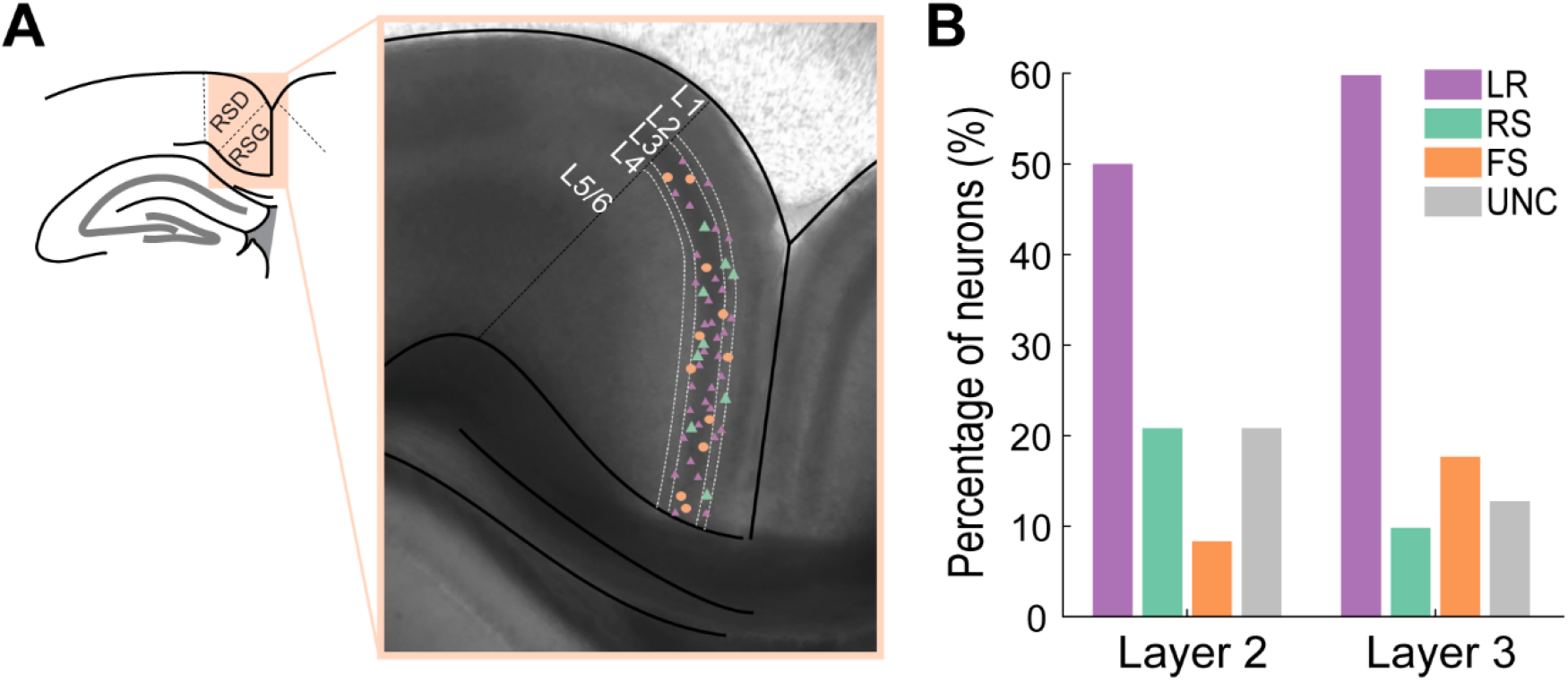
LR cells are the dominant cell type in layers 2/3 of the granular retrosplenial cortex. ***A.*** Representative anatomy of the RSG and locations of a subset of patched neurons. The left panel shows the location of the retrosplenial cortex superior to the corpus callosum. RSD and RSG are distinct regions within the retrosplenial cortex with RSD being more superior and lateral and RSG inferior and medial. Right panel shows a DIC image of a retrosplenial mouse brain slice with RSD and RSG separated by a black dotted line. The layers are demarcated by white dotted lines. Small purple triangles, large green triangles, and orange circles denote representative patch locations of roughly one-half of LR, RS, and FS cells patched in this study, respectively. ***B.*** Percentage of each neuronal subtype patched in each layer of RSG. LR cells are the most prevalent cell type in each layer. UNC stands for unclassified – this group consists of other cell types within this region and neurons that were unable to be classified into one of the other three groups based on their physiological and intrinsic properties.

### Low Rheobase cells are found across the long-axis of the retrosplenial cortex

The retrosplenial cortex is a large structure, spanning 4.38 mm rostracaudally in mice. In addition to LR cells being the most prevalent cell type, we also found that their expression is consistent across this entire long axis of the retrosplenial cortex (Figure 4B). This suggests that the contribution of LR cells to retrosplenial circuit computations is likely to be similar across the long axis of the RSG.

**Figure 4.**
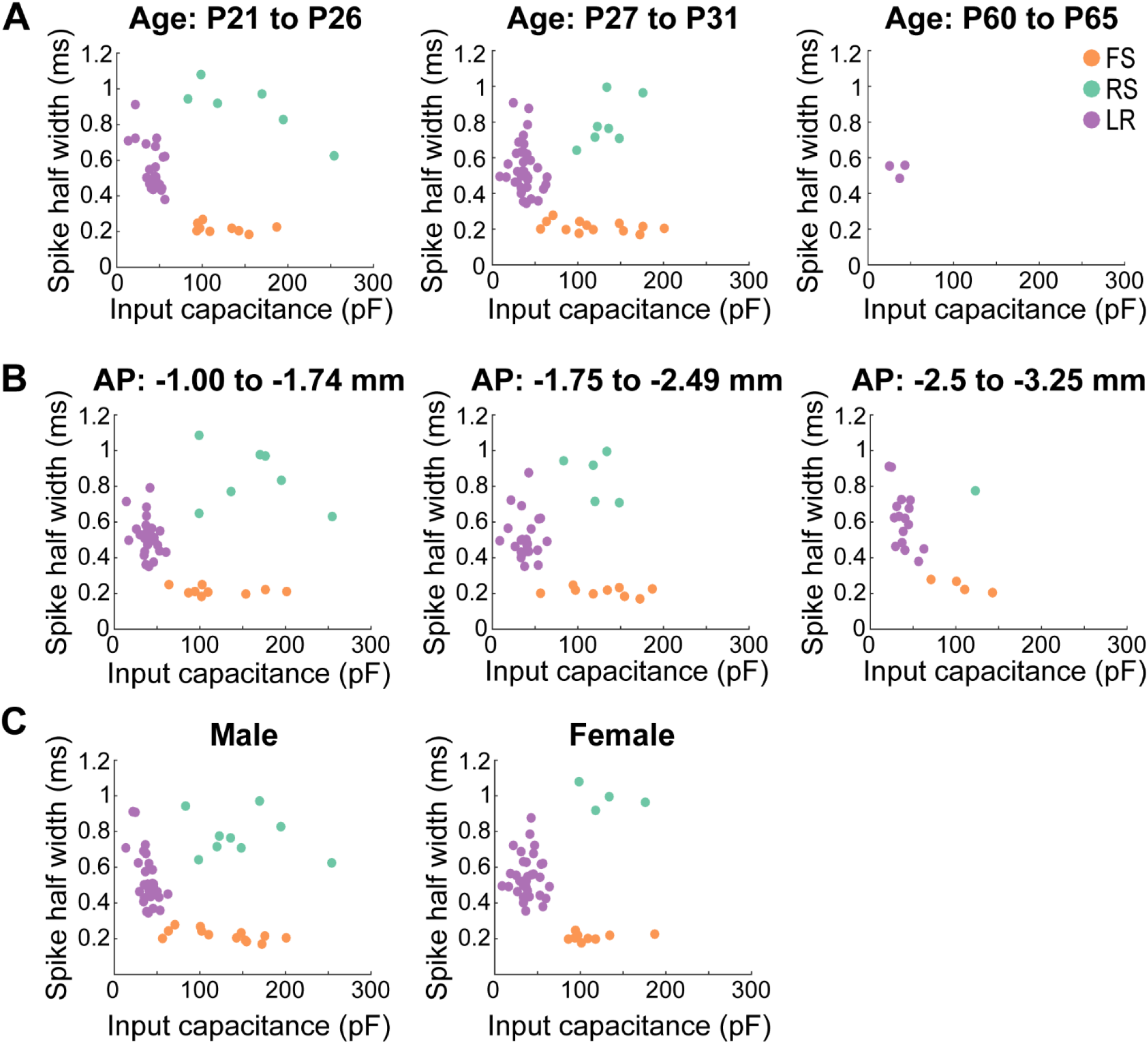
LR cells are consistent across age, sex, and long-axis of the RSG. ***A.*** Scatterplots of the three cell types plotted as a function of spike width and input capacitance across different age groups. Left panel, Postnatal days 21-26 show consistent clustering between the three cell groups (FS: *n* = 9; RS: *n* = 6; LR: *n* = 22). Middle panel, Postnatal days 27-31 show consistent clustering between the three cell groups (FS: *n* = 13; RS: *n* = 7; LR: *n* = 46). Right panel, Postnatal days 60-65 show presence of LR cells in adult mice (*n* = 3) clustering similarly to those recorded in adolescent mice. ***B.*** Similar to ***A***, but now plotted across distinct anterior-posterior sections of the RSG. Left panel, −1 to −1.74 mm from bregma show consistent clustering between the three cell groups (FS: *n* = 9; RS: *n* = 7; LR: *n* = 27). Middle panel, −1.75 to −2.49 mm from bregma show consistent clustering between the three cell groups (FS: *n* = 9; RS: *n* = 5; LR: *n* = 24). Right panel, −2.5 to −3.25 mm from bregma show consistent clustering between the three cell groups (FS: *n* = 4; RS: *n* = 1; LR: *n* = 17). ***C***. Similar to ***A***, but now plotted across sex. All three cell types exist and cluster consistently in both male (FS: *n* = 13; RS: *n* = 9; LR: *n* = 32) and female (FS: *n* = 9; RS: *n* = 4; LR: *n* = 36) mice.

### Low Rheobase cells are found in both males and females at all ages examined

LR cells are present in both adolescent and adult mice, suggesting this highly intrinsically excitable cell is not a transient developmental phenotype (Figure 4A). LR cells are found in both male and female mice (Figure 4C). Thus, these neurons are the dominant cell type in the superficial granular retrosplenial cortex, consistent across age, sex, and long-axis of the RSG.

### Low Rheobase cells are excitatory

In order to investigate whether LR neurons were excitatory or inhibitory, we next conducted whole-cell recordings coupled with optogenetic activation of channelrhodopsin in CaMKII+ cells. CaMKII-Cre x Ai32 mice (Jackson Laboratories 005359 and 024109 respectively, crossed in house) were used for these experiments. In these mice, cells containing the excitatory marker CaMKII express Cre, thus allowing for expression of a cre-dependent channelrhodopsin (ChR2) exclusively in CaMKII neurons (Figure 5A). We then used 1 ms light pulses in a 10 Hz train to test ChR2 responses in the patched neurons. Of 20 LR cells tested, 85% (17/20) directly responded to the optogenetic light pulse, indicating that they were directly expressing ChR2 and thus were CaMKII positive (Figure 5B&C). This suggested, but did not prove, that they may be excitatory neurons.

**Figure 5.**
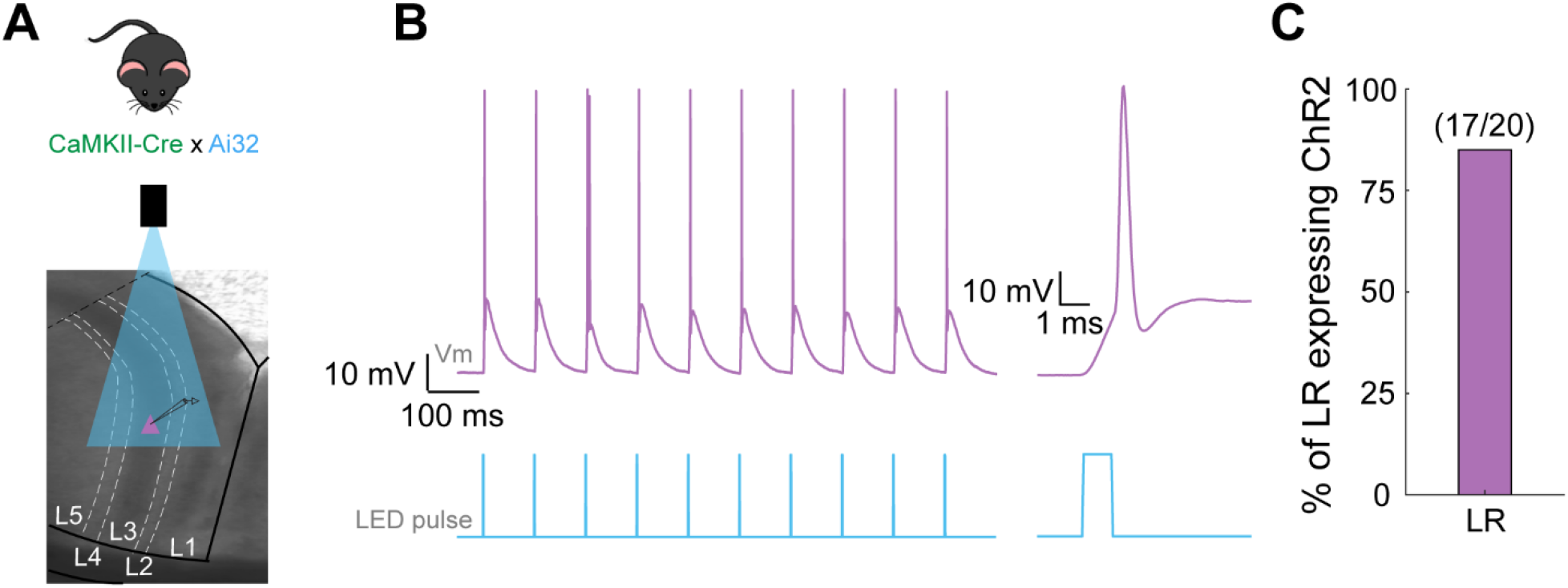
LR cells directly respond to ChR2 in CaMKII-Cre x Ai32 mice, indicating expression of CaMKII in this cell type. ***A.*** Schematic showing the experimental set-up. Top panel, Mouse indicating the genetic cross of CaMKII-Cre (Jackson Laboratories, 005359) and Ai32 (Jackson Laboratories, 024109) on a C57Bl6 background (crossed in house). Bottom panel, Schematic of the experimental set up. 10 Hz optical LED pulses were delivered to layers 2 and 3 of the RSG while the responses of whole-cell patched neurons were recorded. ***B.*** Representative responses of LR cell to the 10 Hz optical pulses. Left traces show all 10 optical pulses (1ms each) over the span of one second and the patched LR cell spiking one to two times in response to each 1ms LED pulse. Right trace is a zoomed in view of the first optical pulse and resulting spike from the LR cell. The almost instantaneous neuronal response to the light (<0.15 ms latency) is indicative of direct ChR2 expression. ***C.*** Bar graph representing the percentage of LR cells tested that directly expressed ChR2 (85%, 17/20).

We then confirmed the excitatory nature of LR cells using paired recordings of layer 2/3 RSG neurons. Although connections were rare, when LR cells were connected to neighboring cells, they led to excitatory post-synaptic potentials (EPSPs) in their paired FS cell (Figure 6D). This is the first demonstration that LR cells in RSG are indeed excitatory neurons.

**Figure 6.**
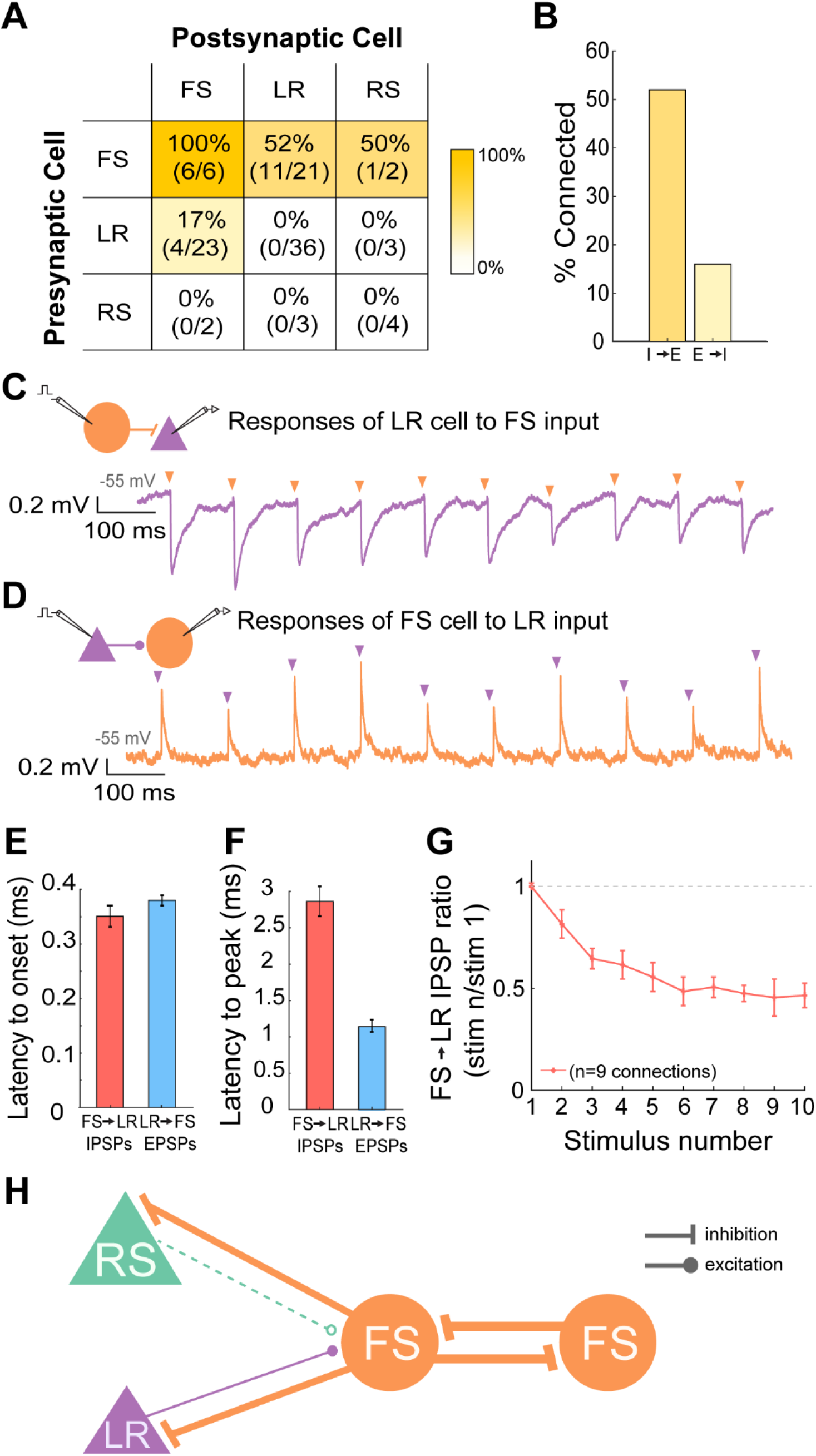
Dominant local inhibition in the superficial layers of the retrosplenial cortex. ***A.*** Table indicating the percentage of connectivity between all types of pairs tested. FS-FS connections were found in all 6 directions tested (100%). FS→LR connections existed in 11 of the 21 pairs recorded (52%). FS→RS connections existed in 1 of the 2 pairs recorded (50%). LR→FS connections existed in 4 of the 23 pairs recorded (17%). There were no RS→FS connections recorded and no E→E connections recorded (LR→LR, LR→RS, RS→LR, RS→RS). The heat map indicates the probability of connections between the neuron types identified in each cell of the table. Deeper gold indicates connection probabilities of near 100%, while lighter gold represents lower probabilities, and white denotes a connection probability of 0. ***B.*** Bar graph representing the total connectivity probability between all inhibitory to excitatory directional pairs (52%) and all excitatory to inhibitory directional pairs (16%). Bootstrap resampling followed by a t-test revealed a significantly higher likelihood of observing I→E connections versus E→I connections. ***C.*** Representative trace of the connection between a presynaptic layer 3 FS cell and a postsynaptic layer 3 LR cell (held at −55 mV). The neurons were 27 um apart with the LR cell located superficial to the FS cell. Schematic shows the patched pair in which the FS cell is being stimulated to spike at 10 Hz, with postsynaptic potentials recorded in the LR cell. The purple trace is the response of the LR cell to a 10 Hz sequence of FS cell spikes (indicated by the orange arrows). ***D.*** Similar to C, but now for a presynaptic LR to postsynaptic FS excitatory connection. ***E.*** Bar graph showing the average latency to onset of the IPSPs recorded from the FS−LR pairs (red) and the EPSPs recorded from the LR−FS pairs (blue) [p=0.9273; Wilcoxon rank sum test]. Error bars are standard error. Latency to onset was calculated as the time from the peak of the presynaptic action potential to the beginning of the postsynaptic IPSP/EPSP. ***F.*** Bar graph showing the average latency to peak of the IPSPs recorded from the FS→LR pairs (red) and the EPSPs recorded from the LR→FS pairs (blue) [p=0.009; Wilcoxon rank sum test]. Error bars are standard error. Latency to peak was calculated as the time from the peak of the presynaptic action potential to the peak of the postsynaptic IPSP/EPSP. ***G.*** Group synaptic dynamics for FS→LR connections (n=9). Inhibition onto LR cells exhibited strong short-term depression. ***H.*** Schematic of the microcircuitry of FS, RS, and LR neurons in the superficial layers of RSG.

### Dominant inhibition and rare local excitation in the superficial layers of RSG

Using paired whole-cell recordings, we sought to quantify the connectivity between these three major cell types in the superficial layers of RSG: LR and RS (both excitatory; E) and FS (the major inhibitory neurons in these layers; I). To our surprise, LR to FS connectivity was rare (17%), suggesting a relative lack of locally driven excitation of FS cells. On the other hand, FS cells were frequently connected to, and inhibited, neighboring LR cells (52%) (Figure 6A). When all pairs were considered, the E→I connectivity was only 16%, whereas the I→E connectivity reached 53% (Figure 6B). The difference in probability to observe I→E connections versus E→I connections was significant (p<0.01; two-tailed t-test), suggesting the superficial layers of the RSG represents an inhibition-dominated network, with feedforward inhibition far more likely than feedback inhibition. Additionally, we observed no LR→LR connections (0/36), nor any connectivity between LR and RS cells (0/6), indicating a complete lack of E→E connectivity. In contrast, FS→FS connectivity was robust, being found in each of the 6 directions tested across three pairs (100%; Figure 6A).

The latency to onset of the evoked responses from a holding potential of −55 mV was similar between inhibition and excitation (p=0.9273; Wilcoxon rank sum test; Figure 6E). However, the peak of the EPSP from LR onto FS cells was reached significantly faster than the peak of IPSPs from FS to LR (p=0.0091; Wilcoxon rank sum Test; N=3 LR→FS connections; N=9 FS LR connections; Figure 6F). IPSPs from FS to LR cells exhibited clear short-term depression. This was seen in paired recordings (Figure 6C&G) and also when recording from LR neurons during optogenetic stimulation of FS cells (data not shown). EPSPs from LR to FS cells did not clearly exhibit either depression or facilitation (Figure 6D). The circuit diagram for L2/3 of RSG is summarized in Figure 6H and highlights the prominent role of inhibition in this circuit.

### Axons from LR cells do not ramify locally but head to deeper layers and towards the corpus callosum

The rarity of connections from LR neurons onto their neighboring L2/3 cells suggested that LR axons have more distant targets. In order to investigate the projections of the LR cells, we used biocytin to fill cells for morphological consideration after characterizing their physiological properties. Of the 3 LR neurons whose cell body, dendrites, and axons were sufficiently filled, all exhibited projections to the deeper layers of RSG (Figure 7A,B,D). Of the three, one axon clearly entered and traveled within the corpus callosum (Figure 7A&C). Additionally, LR neurons had very few axonal ramifications within layers 2/3, matching their extremely low likelihood to synapse onto local neurons (Figure 7A&D). Upon further examination of our four paired recordings in which LR cells directly excited the paired FS cell, we noted that all of these LR cells were located more superficially than their paired FS cell. Conversely, of the four pairs in which the FS cell was located more superficial to the LR cell, none exhibited connections from the LR to the FS cell. This supports the finding that LR axons travel to deeper areas and do not ramify locally or superficial to their cell bodies. It also suggests that FS cells in L2, even more so than L3 FS cells, are likely to be completely devoid of local excitation from LR cells.

**Figure 7.**
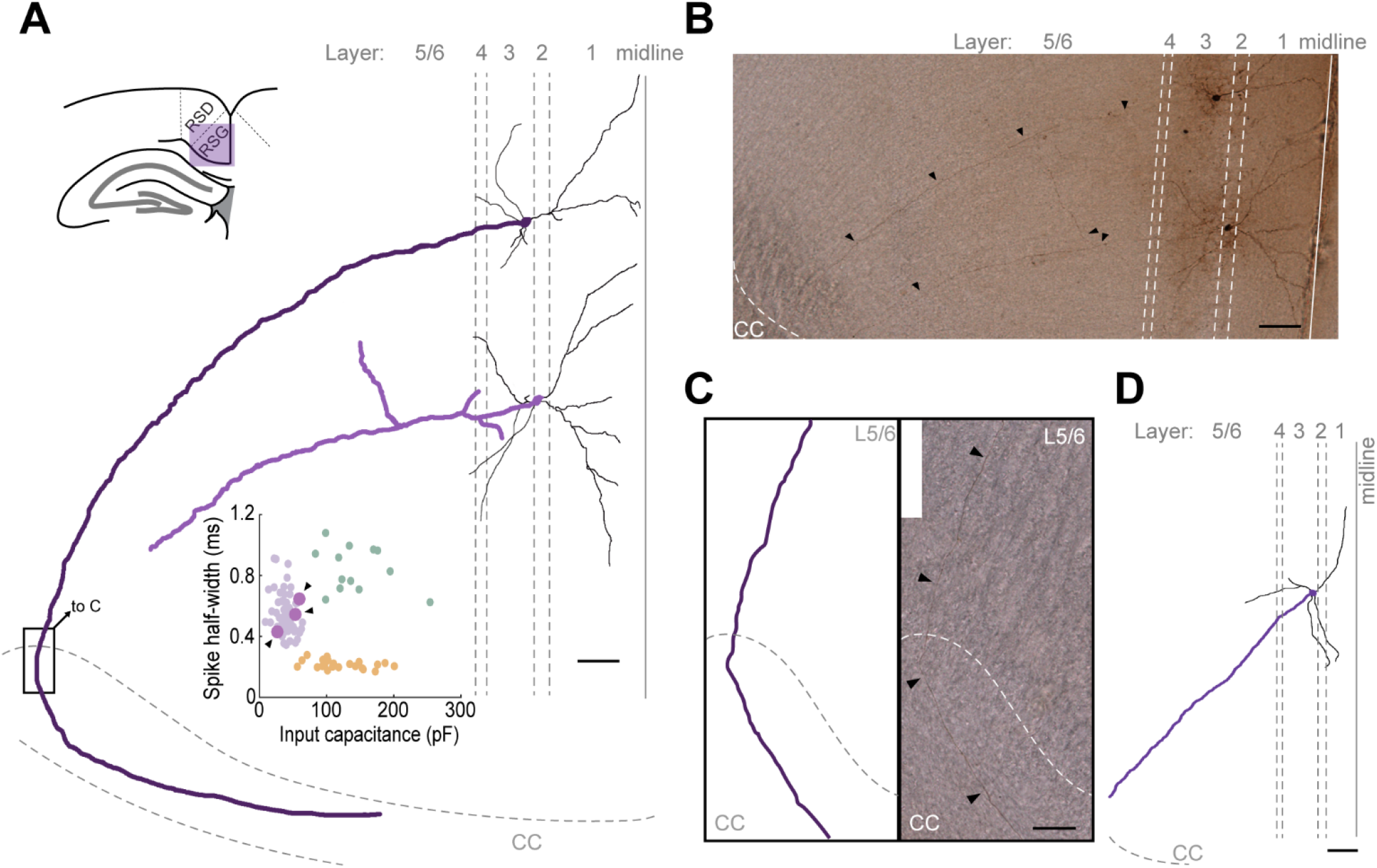
LR axons do not ramify locally and instead project to deeper layers and the corpus callosum. ***A.*** Schematic of the axonal ramifications of two LR neurons whose cell bodies are located within L2/3. Top left shows the location of the RSC within the slice. A slice with an anterior-posterior location of −1.82 mm was used in a P25 mouse. Layers and corpus callosum (CC) are demarcated by grey dashed lines, with the midline of the brain demarcated as a solid line. Scale bar represents 50 um. Dendrites are in black, and cell bodies/axons are in purple. Axons project clearly to deeper layers, sometimes entering the corpus callosum. Minimal axonal ramifications are observed in L2/3. Inset is identical to that in Figure 2C with the three LR cells referenced here in larger purple dots, indicated by the arrows. All three LR cells cluster clearly within the LR cell group and separate from RS and FS cell groups. ***B.*** Biocytin fill used to create the schematic in A. Arrows are placed periodically along the axon for visualization. Scale bar represents 50 um. ***C.*** Zoomed in view of the indicated box in A. Left shows a schematic of the axon projecting through L5/6 before entering and traveling within the CC. Right shows biocytin fill image. Arrows are placed periodically along the axon for visualization. Scale bar represents 50 um. ***D***. Schematic of the third filled LR cell dendrites, cell body, and axon. A slice with an anterior-posterior location of −2.54 mm was used in a P24 mouse. Layers and corpus callosum are demarcated by grey dashed lines; a solid line indicates the midline. There are no axonal ramifications in the superficial layers, and the axon projects into deeper layers towards the corpus callosum. Scale bar represents 100 um.

### LR, but not RS, neurons support high fidelity, sustained responses to persistent head-direction inputs

We next created biophysically-realistic models of both LR and RS cells (see Methods). Multi-compartmental models consisting of a soma and dendritic compartments were used in each case, and the dendritic morphology was matched to the dense dendrites of RSC RS neurons compared to the far fewer branches seen in LR cells (Figure 8A, 8E). Experimental physiological properties were accurately reproduced in each model (Figure 8B, 8F), with the LR neuron model having a higher input resistance, minimal spike frequency adaptation, and narrower spike width than the RS neuron model (Figure 8C, 8G). Frequency-current responses and latency-current response of the neuron models also closely corresponded to the experimental data for each neuronal subtype (Figure 8D, 8H).

**Figure 8.**
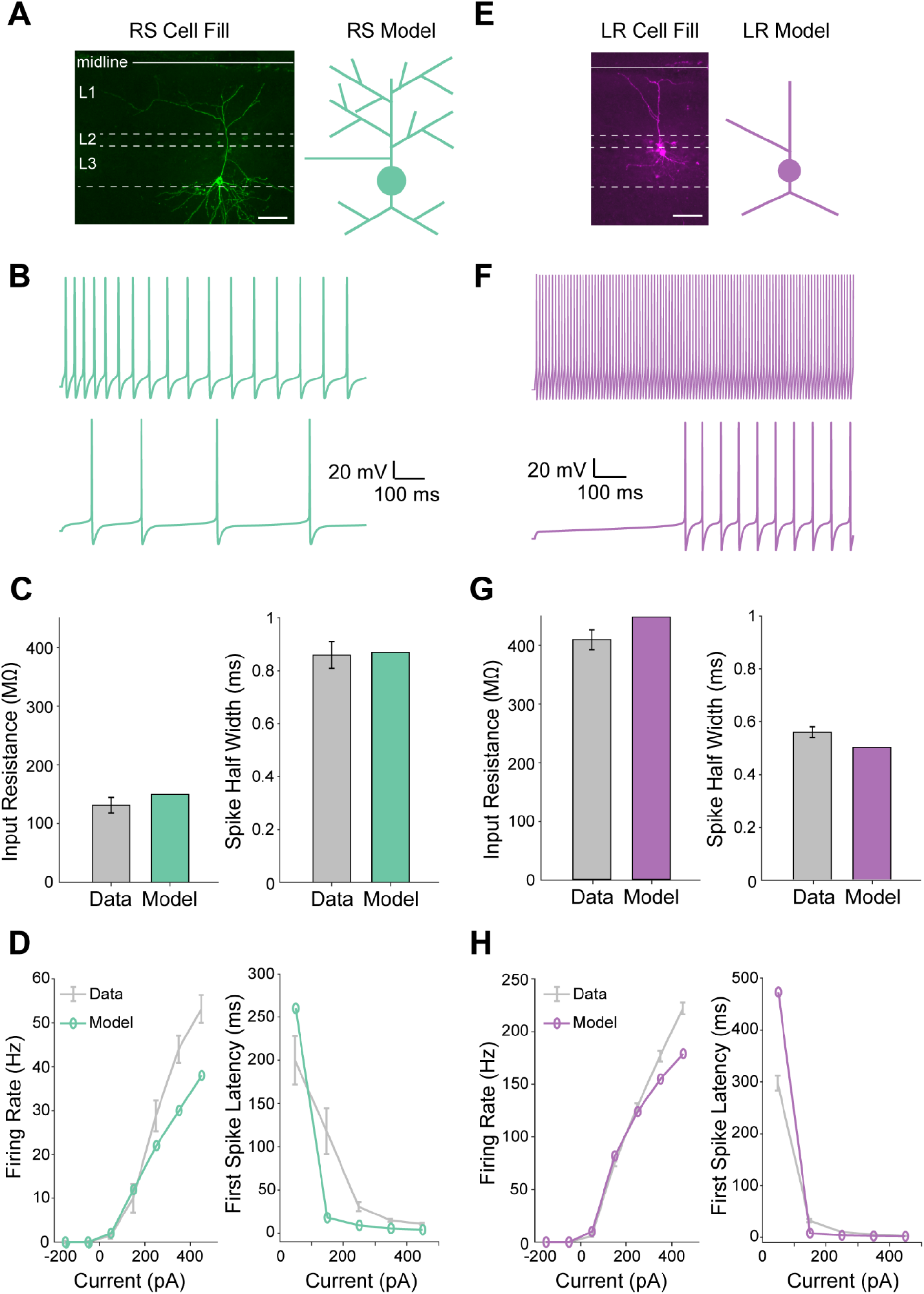
Computational models of LR and RS neurons accurately replicate the physiological and morphological properties of the respective neuronal subtypes. ***A.*** Left, fluorescent fill of a L3 RS cell showing extensive dendritic branching. Scale bar indicates 50 μm. Right, Morphology of the corresponding RS neuron model. ***B.*** Response of the RS neuron model to a low (100 pA) and high (200 pA) current injection. ***C.*** Left, Input resistance of the RS neuron model (green; 150 MΩ) compared to the average input resistance of the physiologically recorded RS cells (grey; 131.27 MΩ). Right, Bar graph representing the spike width of the RS neuron model (green; 0.87 ms) compared to the average spike width of the physiologically recorded RS cells (grey; 0.86 ms). ***D.*** Left, Similar Frequency-Current (F-I) relationships for the model (green) and physiologically recorded (grey) RS cells. Right, Latency to first spike-Current (L-I) relationship for the model (green) and physiologically recorded (grey) RS cells. ***E.*** Left, Fluorescent fill of a L3 LR cell showing minor dendritic branching. Scale bar indicates 50 μm. Right, Morphology of the LR neuron model. ***F.*** Modeled LR cell response to a low (50 pA) and high (200 pA) current injection. ***G.*** Left, Input resistance of the LR neuron model (purple; 441 MΩ) compared to the average input resistance of the physiologically recorded LR cells (grey; 402.69 MΩ). Right, Bar graph representing the spike width of the LR neuron model (purple; 0.49 ms) compared to the average spike width of the physiologically recorded LR cells (grey; 0.55 ms). ***H.*** Left, Similar F-I relationship for the model (purple) and physiologically recorded (grey) LR cells. Right, L-I relationship for the model (purple) and physiologically recorded (grey) LR cells.

We next used these biophysically-realistic models to understand the information processing capabilities of LR versus RS neurons. The subicular complex provides one of the most prominent sources of input to the superficial layers of the RSC (Yamawaki et al., 2019a). Most cells in the subicular complex fire selectively when an animal is facing a particular direction and are hence called head-direction (HD) cells (Taube et al., 1990). These subicular HD cells are bursty, firing a series of rapid spikes at a rate of 150-250 Hz (Simonnet and Brecht, 2019). The same subicular HD cells also fire long-duration, persistent trains of action potentials (Taube and Bassett, 2003; Yoshida and Hasselmo, 2009; Peyrache et al., 2015). This persistent firing is thought to be critical for maintaining a sense of orientation when the animal is not moving, but instead continuously facing a particular direction (Taube and Bassett, 2003; Yoshida and Hasselmo, 2009; Peyrache et al., 2015).

We sequentially examined how these important properties of subicular inputs are processed by RS versus LR neurons. First, 200 Hz bursts consisting of 5 spikes were input into both RS and LR cells (Figure 9A, 9B). The RS cell response to each of the constituent spikes within the burst was characterized by a low probability of firing and imprecisely timed action potentials (high jitter). LR cells, on the other hand, responded with high reliability and more precisely timed action potentials with little jitter across trials (Figures 9C-9F). LR cell spikes were significantly more reliable and more precise (less jitter) than RS cell spikes in response to the burst input (p<0.001; two-tailed t-test), showing that LR neurons are capable of higher fidelity burst encoding with superior spike timing coding capabilities than their neighboring RS neurons. Qualitatively similar results were obtained when the Gmax of synaptic inputs was halved in strength (data not shown).

**Figure 9.**
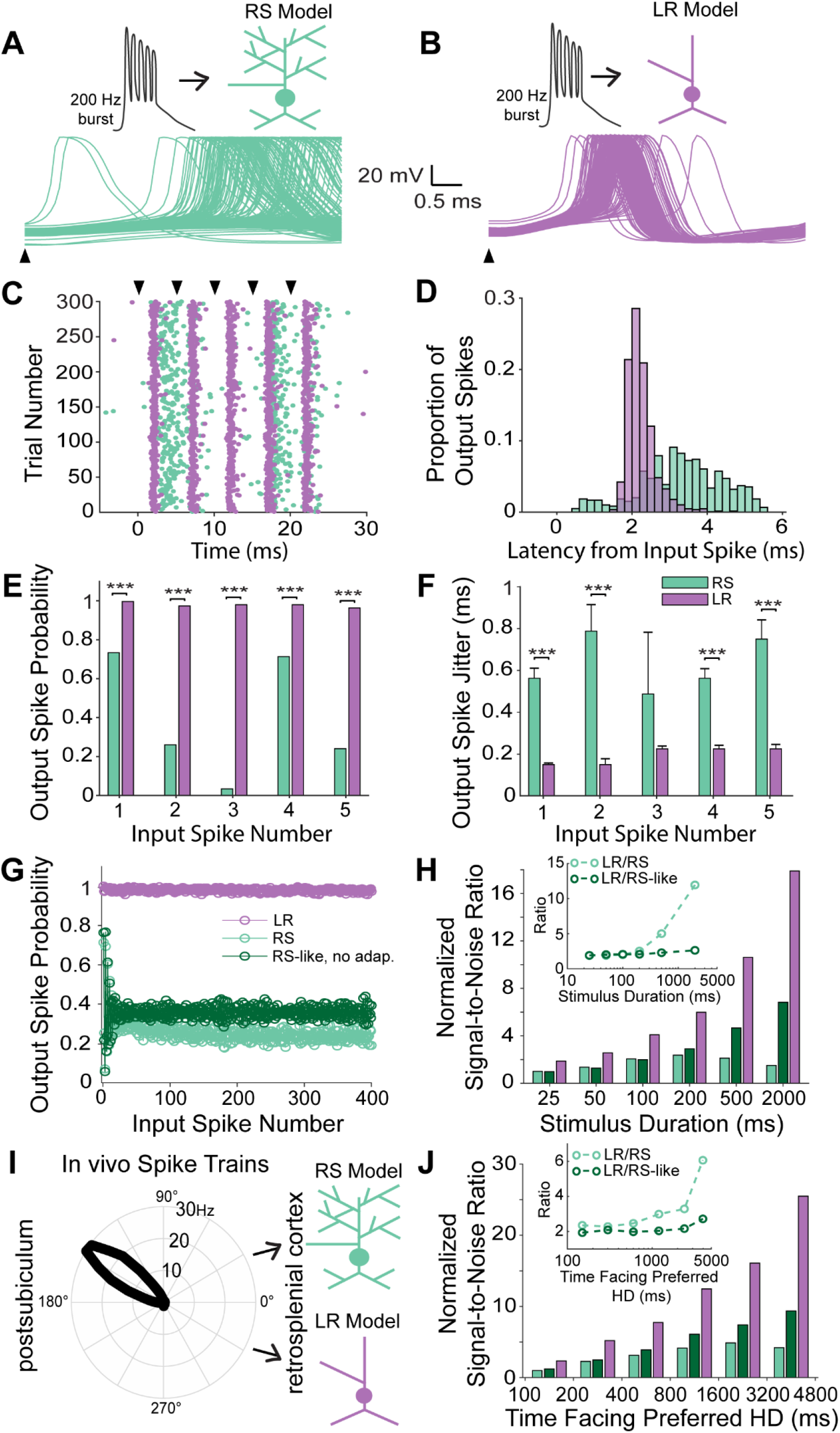
Unique properties of LR neurons enable high fidelity, sustained transmission of persistent head-direction input. ***A.*** Top, Schematic of a 200 Hz, 5-spike burst input to the RS neuron model. Bottom, Membrane potential traces of the RS neuron in response to first spike of the burst input across 300 trials. ***B.*** Top, Similar to ***A***, but for the LR neuron model. ***C.*** Raster plot of the firing of RS and LR neuron model in response to the burst input. RS neuron model response is characterized by low-reliability and lack of spike timing precision, whereas LR neurons fire with high reliability and high spike timing precision. ***D.*** Distribution of spike latencies of the RS (green) and LR (purple) neuron model for all five spikes within the burst input, highlight the precise timing of LR responses. ***E.*** Output spike probability of the RS (green) and LR (purple) neuron model to each individual spike within the burst. LR model has a significantly higher probability to respond to each spike within the burst compared to RS neuron model [*** p<0.001; Pearson chi-square test]. ***F.*** Spike timing precision computed as the jitter in the output spikes for the RS (green) and LR (purple) neuron model to each individual spike within the burst. LR neuron model is characterized by high spike timing precision compared to RS cells [*** p<0.001; two-tailed t-test]. ***G.*** Output spike probability of the LR model and two models of RS cells (a standard model and a second model without adaptation) as a function of input spike number for a 2 second continuous stimulation. Note the diminished spike probability on both RS models compared to the highly reliable LR responses. ***H.*** Signal to noise ratio (SNR) of RS (with and without spike frequency adaptation) and LR neuron models as a function of the duration of the input stimulus. Inset represents the ratio of SNR of the LR neuron model to the SNR of the RS neuron model (with and without spike frequency adaptation). Difference in SNR between RS and LR neuron increases disproportionately for larger input durations due to the effect of the RS neuron’s spike frequency adaptation. Hence, RS neurons lose the ability to distinguish their spiking response to sustained signals from background noise under conditions of continuous input. ***I.*** Schematic of *in vivo* head direction input from the postsubiculum (from dataset: Peyrache, A., Buzsáki, G. (2015)) to the modeled RS and LR neurons. The polar plot shows the *in vivo* head-direction tuning of the input postsubiculum head-direction cell. ***J.*** Same as H, but now in response to realistic *in vivo* spike trains of the postsubiculum head-direction cell shown in ***I***. Similar to continuous burst stimulation, the RS neuron model’s SNR declines rapidly with increase in stimulus duration due to spike frequency adaptation, highlighting the ability of LR, but not RS, neurons to respond to persistent head-direction input.

We next examined the ability of LR and RS neurons to respond to persistent inputs of varying durations. First, we utilized a continuous input spike train of 200 Hz over progressively longer durations. LR cells were able to respond with high probability and high precision for all durations examined (Figure 9G). RS cells, on the other hand, had average response probabilities under 0.4, even when we removed the adaptation current from the RS cell model. Signal-to-noise ratio (SNR) analyses (see Methods) showed that the LR cell response was characterized by high SNR to prolonged 200 Hz inputs (Figure 9H). RS cells, with and without adaptation, showed several fold lower SNR for both short and long durations (Figure 9H). The ratio of SNR_LR_ to SNR_RS_ progressively increased at higher input durations (Figure 9H, inset), indicating the progressive failure of RS cells to reliably encode longer duration inputs with enough spikes. When we removed the adaptation current from the RS cell model, the ratio SNR_LR_ to SNR_RS-No-Adap_ remained very high but did not progressively increase as a function of increasing input durations (Figure 9H, inset), indicating that the spike frequency adaptation of RS neurons prevents reliable transmission of long duration, persistent inputs to its postsynaptic targets. These results suggest that a combination of passive and active properties enable LR neurons to have higher fidelity and sustained SNR at high input durations, whereas the spike frequency adaptation of RS neurons amplifies the SNR disadvantage of RS neurons at longer durations.

Finally, we validated our models’ predictions by utilizing realistic input spike trains recorded from HD cells in the postsubiculum (Figure 9I; Peyrache et al., 2015; Peyrache and Buzsáki, 2015). As expected, this dataset included epochs of persistent, long-duration firing when the animal faced a given cell’s preferred head direction. We found that LR neurons encoded this postsubiculum HD input with higher SNR at all input durations. The ratio of SNR_LR_ to SNR_RS_ again increased dramatically at higher input durations (when the animal faced the same direction for long periods of time; Figure 9J), indicating the inability of RS neurons to faithfully encode persistent, long duration inputs coming from the subicular HD cells. Thus, the intrinsic properties of LR neurons mean that they are better suited to encode persistent HD inputs than RS cells.

## DISCUSSION

The unique cytoarchitecture of the retrosplenial cortex has long been appreciated by neuroanatomists (Rose, 1927; Wyss et al., 1990; Wyss and van Groen, 1992; Ichinohe and Rockland, 2002; van Groen and Wyss, 2003). The granular division of the retrosplenial cortex, in particular, has two geometric features that appear to set it apart from many other cortical regions: 1) uniquely small pyramidal neurons that cluster most densely in layers 2 and 3 (Wyss et al., 1990; Kurotani et al., 2013); 2) the bundling of apical dendrites emanating from these L2/3 pyramidal neurons (Wyss et al., 1990; Ichinohe and Rockland, 2002). It is thus not a great leap to suggest that a thorough understanding of granular RSC function would greatly benefit from an in depth understanding of these unique L2/3 neurons.

Here, we have characterized the detailed intrinsic properties, connectivity, and computational capabilities of these small pyramidal cells, as well as their neighboring, larger, more familiar RS cells, revealing a number of key differences. We call these small pyramidal cells “Low Rheobase” (LR) neurons based upon the ease with which they can be excited and made to fire (due, in large part, to their high input resistance). In addition to their high excitability, these cells also have spike widths that are much narrower than RS cells and show minimal spike frequency adaptation, again in sharp contrast to RS neurons (Figure 1; Table 1). We also found that all three cell types examined had a tendency to spike late during a near threshold current injection, consistent with high levels of Kv1.1 or Kv1.2 ion channel expression in the RSC (Kurotani et al., 2013). However, this was most pronounced in FS cells, and not in LR cells (Figure 1). Furthermore, RS cells also showed a milder late spiking phenotype (Figure 1). Thus, we believe the key, unique computational features of LR cells are their hyperexcitability and lack of spike-frequency adaptation.

How do the distinct passive and active properties of LR versus RS neurons impact their input-output transformations and information-coding capabilities? We used biophysically-realistic models to investigate this question. On the short time-scale, the higher excitability, shorter spike widths, and more electrotonically compact dendritic tree of LR cells enables them to spike with a higher probability and lower jitter in response to incoming bursts of spikes (Figure 9A-F), such as those generated by afferent subicular cells during active behaviors (Simonnet and Brecht, 2019). On the long time-scale, the almost complete lack of spike-frequency adaptation helps LR cells maintain sustained (and still precise) responses to incoming persistent inputs (Figure 9G-J). This encoding of persistent information appears critical to the function of the RSC. Recent imaging evidence suggests that RSC neurons (irrespective of subtype) have a unique ability to encode long-duration, history-dependent value signals (Hattori et al., 2019). Of even more direct relevance is the persistent nature of the navigational information being received and processed by the superficial RSC. The subiculum represents one of these key functional inputs to RSC L2/3 cells (Wyss and van Groen, 1992; Yamawaki et al., 2019b, 2019a). Subicular neurons, particularly those in the postsubiculum, display a strong preference for particular orientations and are thus called head direction (HD) cells (Taube et al., 1990). When an animal faces a particular direction for long durations, postsubiculum HD cells keep spiking persistently, likely contributing to the maintenance of the sense of orientation in the absence of ongoing vestibular changes (Taube et al., 1990; Chrastil et al., 2017). The unique properties of LR cells suggest they are ideally suited to respond to this persistent subicular input (Figure 9I-J), enabling the RSC to utilize this valuable head-direction input to help generate a sense of orientation regardless of how long an animal has been facing the same direction. In fact, our results suggest that the longer an animal faces a postsubicular cell’s preferred direction, the better the signal-to-noise ratio with which LR neurons can encode this information (due to the accumulation of high rate signal spikes over time), potentially helping to increase the behavioral certainty of the current orientation and helping to recall orientation-relevant memories. Indeed, a sense of spatial disorientation is one of the key deficits after RSC damage in humans (Bottini et al., 1990; Takahashi et al., 1997; Ino et al., 2007; Osawa et al., 2007).

Several lines of future work are needed to better understand LR versus RS neuronal processing in support of RSC function. It is not yet known whether distinct types of subicular cells (Simonnet and Brecht, 2019; Yamawaki et al., 2019a) differentially contact LR versus RS cells, nor do we yet know the short-term dynamics of subicular inputs to LR versus RS cells. Similarly, it remains to be determined whether LR and RS neurons can independently generate sustained firing in the absence of subicular input: do either LR or RS neurons express the same ion channels that allow postsubicular cells to spike persistently (Chrastil et al., 2017)? Perhaps the single most important gap in the field’s knowledge regarding L2/3 RSC neurons is the lack of any information on their *in vivo* spike patterns and how their firing encodes navigation and memory related information. To our knowledge, no *in vivo* awake, behavioral recordings to date have specifically attempted to precisely target L2/3 of the granular RSC, likely due to its relatively inaccessible position at the midline. Our precise characterization of differences in spike width and adaptation between LR and RS neurons will help the field to identify these distinct neurons using precisely planned extracellular recordings.

The presence of two distinct, neighboring principal neurons is seen in several structures that are important for spatial navigation and memory. Recent work has shown that granule and mossy cells in the dentate gyrus differentially encode spatial information (Scharfman, 1992; GoodSmith et al., 2017; Senzai and Buzsáki, 2017), resolving several previously confusing data points regarding the nature of the sparse code used by granule cells (Leutgeb et al., 2007). Similarly, neighboring deep versus superficial CA1 pyramidal cells have been shown to have differential spatial and temporal firing properties, as well as distinct local and distant connectivity (Mizuseki et al., 2011; Lee et al., 2014; Danielson et al., 2016; Soltesz and Losonczy, 2018). In the subiculum, the very cells that are providing inputs to LR and RS neurons in the RSC are themselves diverse, showing two distinct patterns of bursting as well as differences in VGlut1 versus VGlut2 expression (Simonnet and Brecht, 2019; Yamawaki et al., 2019a). Thus, the notion of parallel coding schemes implemented by distinct populations of principal neurons is of clear importance in regions involved in memory and navigation. In the superficial RSC, our results show that such parallel neural codes are likely to be implemented by the distinct properties of LR and RS neurons.

Our paired recordings provide the first direct proof that LR neurons are indeed excitatory (Figure 6). Although LR cells are the most prevalent cell type in L2/3 (Figure 3), they make few local connections (Figure 6), instead sending their axons into the deeper layers and corpus callosum (Figure 7). LR cells receive prominent inhibitory inputs from the neighboring L2/3 FS cells (Figure 6), with the probability of FS-to-LR connectivity reaching 52%, somewhat higher than that reported in many other regions of the neocortex (Beierlein et al., 2003; Yoshimura and Callaway, 2005; Packer and Yuste, 2011; Jiang et al., 2015). This, coupled with the complete lack of local excitatory connections onto LR cells (Figure 6) and the dense FS-FS connectivity (Figure 6), indicates that the superficial layers of the RSG are a network dominated by local inhibition. The inhibition from FS to LR neurons showed similar short-term depression to that seen from FS to RS cells in many other cortical structures (Figure 6G; Beierlein et al., 2003). While we did not explicitly model feedforward inhibition in our simulations, this depression is likely to further aid in the long-duration firing of LR neurons in response to persistent subicular inputs by curtailing the strength of feedforward inhibition over time. Of importance to network computations, the strong FS-FS and FS-LR connectivity is also likely to allow the RSC circuit to implement high frequency oscillations that are generated by the interneuron-gamma (ING) mechanism instead of, or in addition to, oscillations generated via the pyramidal-interneuron-gamma (PING) mechanism. Future large-scale circuit models of the superficial RSC incorporating LR, RS, and FS cells based on the intrinsic properties and connectivity principles described here will help to understand the network computations performed by this unique retrosplenial circuit (Figure 6H).

## METHODS

### Physiological Experimental Methods

#### Slice preparation

All housing of animals and procedures were approved by the University of Michigan Institutional Animal Care and Use Committee. Multiple mouse lines were used in this study, including PV-IRES-Cre (Jackson Laboratories, 008069), CaMKII-Cre (Jackson Laboratories, 005359), Ai32 (Jackson Laboratories, 024109), Ai14 (Jackson Laboratories, 007914), PV-IRES-Cre x Ai14 (crossed in house), PV-IRES-Cre x Ai32 (crossed in house), CaMKII-Cre x Ai32 (crossed in house), and NTSR1-Cre (MMRRC, 030648-UCD). A total of 167 recordings are included in this study from the following mouse lines: PV-IRES-Cre (55), CaMKII-Cre (15), Ai32 (3), PV-IRES-Cre x Ai14 (3), PV-IRES-Cre x Ai32 (64), CaMKII-Cre x Ai32 (20), and NTSR1-Cre (7). Mice of both sexes between the ages of P21-31 and P60-65 were included in the experiments. No differences in cell type properties were observed across mouse lines, while both age and sex were explicitly analyzed in terms of their relationship to cell type properties (see Results).

Mice underwent deep isoflurane anesthesia before decapitation. Brains were removed within one minute of decapitation and placed in an ice-cold high-sucrose slicing solution that had been saturated with carbogen gas for at least 30 minutes prior to use. Coronal slices (300um) were cut using a Leica 1200 VT vibratome. Slices were allowed to rest in the slicing solution for about 2 minutes before being placed in a carbogen-saturated high-magnesium artificial CSF (ACSF) solution to incubate at body temperature (32°C) for 20 minutes. The entire bubbling bath was then removed from the heater, allowing the slices to gradually cool to room temperature. Slices rested an additional 30 minutes at room temperature before use.

Slices were submerged in a recording chamber with a constant flow of ACSF containing 126 mM NaCl, 1.25 mM NaH2PO4, 26 mM NaHCO3, 3 mM KCl, 10 mM dextrose, 1.20 mM CaCl2, and 1 mM MgSO4. Recordings were done between 29-31°C with an ACSF flow rate of 2 mL per minute. All recordings were done within 8 hours of slicing to ensure reputable health of the cells. Patch pipettes with 2-3 um diameter and resistances of 3-6 MΩ were filled with a potassium gluconate internal solution containing 130 mM K-gluconate, 2 mM NaCl, 4 mM KCl, 10 mM HEPES, 0.2 mM EGTA, 0.3 mM GTP-Tris, 14 mM phosphocreatine-Tris, and 4 mM ATP-Mg (pH 7.25, ~290 mOsm).

#### Whole-cell recordings, analysis, & statistics

Slices were visualized using an Olympus BX51WI microscope equipped with Olympus 5x and 60x water immersion lens and the Andor Neo sCMOS camera (Oxford Instruments, Abingdon, Oxfordshire, UK). In most cases, neurons were patched randomly within layers 2/3 of RSG with the exception of experiments in which PV neurons were targeted for patching based on their expression of either an eYFP tag (PV-IRES-Cre x Ai32 cross) or a tdTomato tag (PV-IRES-CRE x Ai14 cross). All recordings were done under current clamp conditions using the Multiclamp 700B and Digidata 1440A (Molecular Devices). Neurons were adjusted for series resistances and held at a resting potential of −65 mV (unless otherwise stated) using a constant holding current injection. Recordings were not corrected posthoc for liquid junction potential. In order to characterize the different neuron types, intrinsic and firing properties of recorded neurons were calculated using the Clampfit and Matlab software packages.

The following intrinsic neuronal properties were calculated: resting membrane potential, spike threshold, spike amplitude, spike width, input resistance (R_in_), membrane time constant (*τ*_m_), input capacitance (C_in_), afterhyperpolarization (AHP) amplitude, AHP latency, spike frequency adaptation ratio, and rheobase. Resting membrane potential was recorded within 2 minutes of break-in. Cells with severely depolarized break-in potentials (> −55 mV) were not included in this study. Spike threshold, amplitude, width, AHP amplitude, and AHP latency were calculated by averaging all spikes in the first sweep of a 600 ms current step protocol that elicited a firing rate of at least 5 Hz. Spike threshold is calculated from the peak of the third derivative of membrane potential (Cruikshank et al., 2012). Spike amplitude was measured as the voltage change from the spike threshold to the peak of the action potential. Spike width was calculated as the full-width at half-max of the spike amplitude. AHP amplitude was calculated as the voltage change from spike threshold to the peak negativity of the AHP, and AHP latency as the time from peak of the spike to peak negativity of the AHP. Input resistance (R_in_), membrane time constant (*τ*_m_), and input capacitance (C_in_) were calculated from a series of small negative current steps ranging from −5 pA to −30 pA, creating a deflection in was calculated using Ohm’s law, as the mean membrane potential of −2 to −4 mV. R_in_ voltage change divided by mean current amplitude. *τ*_m_ was calculated by fitting a single exponential to the average of the initial 60 ms voltage response, ignoring the first 20 ms. Cin was then calculated from those two parameters using the formula *τ*_m_ = R_in_×C_in_. Spike frequency adaptation ratio was calculated from the first sweep of the 600ms current step protocol that elicited a firing rate of at least 10Hz (6 spikes per 600ms) using the equation ISI_last_ / ISI_first_. Rheobase was calculated from 1 sec current pulses increasing in steps of 1-5 pA as the minimum current required to elicit at least one action potential.

A two-tailed Wilcoxon rank sum was used to compute the statistical significance between the intrinsic properties of various neuronal subtypes. To establish the statistical significance between the probability of E→I and I→E connections, a bootstrap resampling (1000 bootstrap samples) method was used to generate a distribution of connectivity probabilities (Sudhakar et al., 2017). Statistical significance was then computed using two-tailed t-test with a confidence interval of 95%.

Synaptic connections between neurons were tested using paired whole-cell recordings. 1 ms current pulses were delivered to the presynaptic neuron at 10 Hz for a total of 1 second (10 pulses). The synaptic responses of the postsynaptic neuron were simultaneously recorded, while holding the postsynaptic cell at −55 mV. Latency to onset of an IPSP or EPSP was calculated as the time from the peak of the presynaptic action potential to the onset of the postsynaptic IPSP or EPSP. Latency to peak was calculated as the time from the peak of the presynaptic action potential to the peak of the postsynaptic IPSP or EPSP.

#### Optogenetic testing of CaMKII expression

Optogenetic verification of CaMKII expression was conducted using CaMKII-Cre x Ai32 mice (Jackson Laboratories 005359 and 024109 respectively, crossed in house) in which channelrhodopsin is expressed in CaMKII-Cre-expressing neurons. Slices were visualized with the Olympus BX51WI equipped with Olympus 5x and 60x water immersion lens. Expression of channelrhodopsin was marked by fluorescence of the eYFP tag. Neurons were recorded in the same manner as described above with at least one additional protocol to verify functional expression of the channelrhodopsin. One millisecond optogenetic light pulses with a 5,500K white LED (Mightex; 14.47 mW) were delivered at 10 Hz while the neuronal responses were recorded. Direct expression was verified by responses to the light pulses under 0.15 ms.

#### Morphological investigations with biocytin

To determine patched cells’ morphology, 5mg/ml of biocytin was added to the internal solution of recording electrodes. Cells were filled with biocytin (Sigma, cat. no. B4261) throughout the recording session, and the pipette was left attached to the cell for at least 20 min. At the end of the recording, cells were “zapped” with fifteen 1 Hz pulses of 3-4 nA current to improve the diffusion of biocytin into the axon (Jiang et al., 2015). Slices were left to recover in the recording chamber for 30 min before further processing. A detailed description of the biocytin labelling and processing is available elsewhere (Marx et al., 2012). Briefly, slices were filled with biocytin as described above, placed in 4% paraformaldehyde (PFA; Acros Organics, cat no. B0144942) for 12-15 hours, and then transferred to phosphate buffer solution (PBS). After 24-48 hours in the PBS, slices were incubated in avidin-biocytin (ABC Elite kit, VectaShield) for 12 hours and then treated with peroxidase to reveal cell morphology. Finally, slices were mounted on microscope slides with Mowiol-based embedding medium and allowed to dry for at least 12 hours. Cells were visualized using a Leica DM4000B light microscope equipped with a Leica DMC 6200 CMOS camera.

#### Morphological investigations with Alexa-Fluor 488

To investigate cell morphology using fluorescence, biocytin (5mg/ml, Sigma cat no. B4261) was added to the internal solution. Cells were filled with biocytin for a total of 20-30 min each, and the slices were then moved to 4% PFA (Fisher Scientific, cat no. 50-980-494) for overnight incubation. Afterwards, the slices were washed in PBS, permeabilized in 0.2% Triton-X (Sigma, X-100), and incubated for 48 hours in streptavidin conjugated Alexa Fluor 488 (1mg/1ml diluted to 1:1000, Thermo Fisher Scientific S11223). Slices were mounted on glass slides using Fluoromout-G mounting medium (SouthernBiotech, cat no. 0100-01) and glass coverslips and visualized with Leica 6000B microscope equipped with a 10x objective and QImaging Retiga-SRV Fast 1394 camera.

### Computational Modeling Methods

#### Model motivation

Biophysical modeling was utilized to study in detail the computational properties of LR and RS neurons and the possible coding mechanisms by which they could contribute to the spatial navigation functions of the RSC. To this end, we constructed multi-compartmental, biophysically realistic models of the two neuronal subtypes and tuned the model parameters so that their intrinsic properties closely match their experimental counterparts. The following section explains in detail the active and passive properties of the LR and RS neuron models. Unlike the experimental data, where junction potential was not adjusted for, all membrane potential values listed below for the computational model should be considered adjusted for the junction potential.

### LR neuron model

#### Morphology and passive properties

The LR neuron model consists of a somatic compartment and 7 dendritic compartments. The model has an input resistance and input capacitance of 441 MΩ and 32.4 pF, respectively, closely matching the experimental values of LR neurons. The membrane time constant of the model is 14.29 ms. The maximum distance from the soma to the apical dendritic tip in the model is 177 μm, closely replicating the distance from the cell body of these neurons to the pia. The resting membrane potential of the model is −78.36 mV.

Both LR and RS neuron models were simulated at a temperature of 30°C, and a q10 value of 3 (Bertil Hille, 2001) was used to scale the temperature dependence of ion channel kinetics. The number of segments in each compartment was calculated using the d-lambda rule (Hines and Carnevale, 2001). The axial resistivity for both models is 200 Ω-cm (Vierling-Claassen et al., 2010).

#### Active properties

Three voltage-gated ion channels were simulated for the LR neuron model: Fast sodium current(*I*_*Na*_), delayed rectifier potassium current (*I*_*kdr*_), and K_v_1 current (*I*_*d*_). In addition, a phenomenological mechanism (*I*_*adap*_) for spike frequency adaptation was also modeled (Treves, 1993; Fuhrmann et al., 2017). The properties of these currents are described in detail in the following sections. The LR neuron model has a spike half-width of 0.49 ms and a spike threshold of −52 mV. The model exhibits very little spike frequency adaptation, with an adaptation ratio of 1.1, as seen in the experimental data. For both models, the reversal potential of sodium (*E*_*Na*_) and potassium ions (*E*_*K*_) were set to +50 mV and −96 mV, respectively.

##### Fast sodium current

The fast sodium current (*I*_*Na*_) responsible for action potential generation was modeled based on Hodgkin Huxley formulation (Hodgkin and Huxley, 1952) using the experimental gating properties of transient sodium current found in RS neurons (Martina and Jonas, 1997). The channel was modeled with 3 activation gates and an inactivation gate. The channel was distributed in all the 8 compartments of the model, and their respective maximal channel conductance (*g*_*max*_) is tabulated in Table 2. The channel equations and parameters (Martina and Jonas, 1997) (voltage dependence of steady state activation/inactivation (*m*_∞_,*h*_∞_) the time constants of activation and inactivation (*m*_∞_, *h*_∞_), the time constants of activation and inactivation gates (*τ*_*m*_,*τ*_*h*_), channel current (*I*_*Na*_)) is given below.

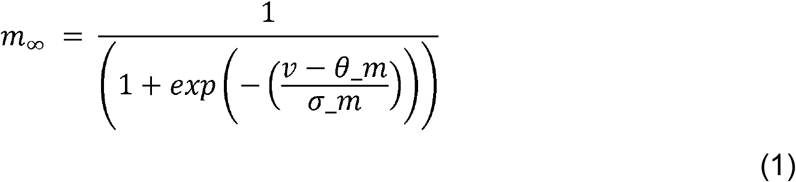

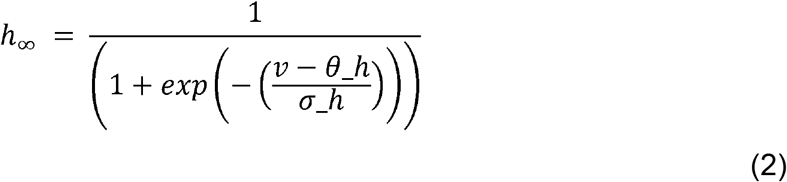

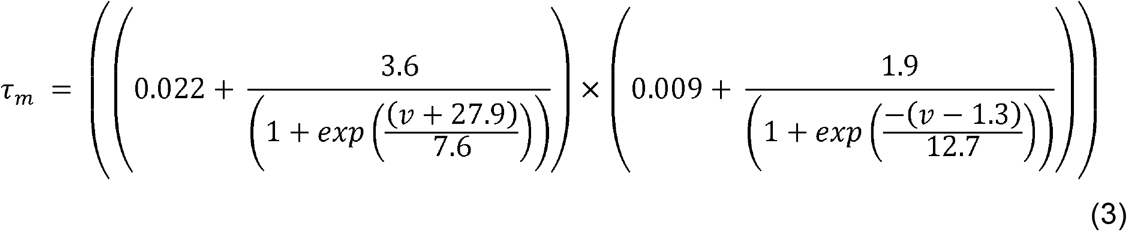

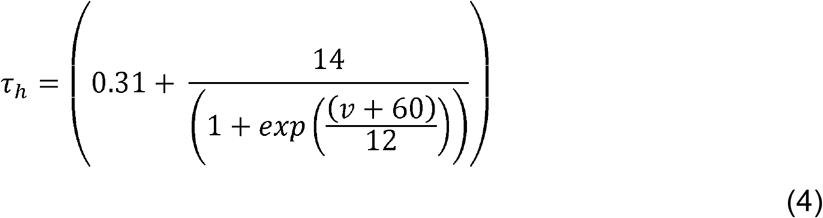

where *θ*_*m*_ = −22.8 *mV*, *σ*_*m*_ = 11.8 *mV*, *θ*_*h*_ = −62.9 *mV*, *σ*_*h*_ = −10.7 *mV*.

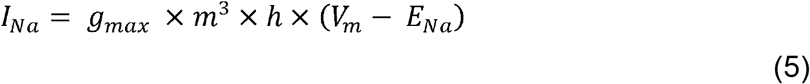

**Table 2.**
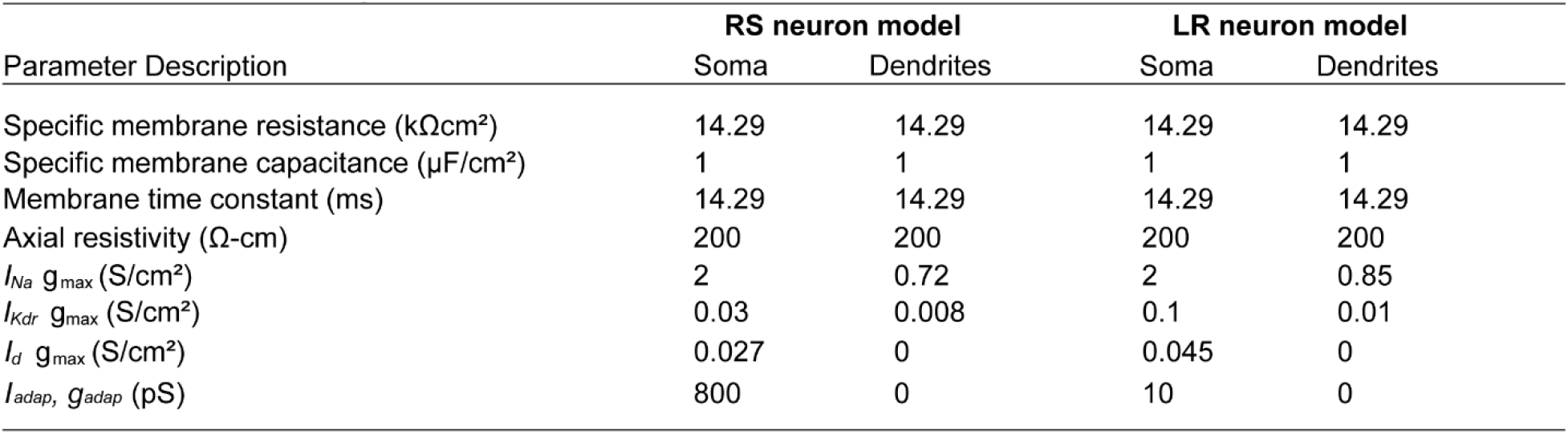
Model parameters. The table lists the values of various model parameters and distribution of ion channel conductances in the somatic and dendritic compartments of the LR and RS neuron models.

| Parameter Description | RS neuron model |  | LR neuron model |  |
| --- | --- | --- | --- | --- |
|  | Soma | Dendrites | Soma | Dendrites |
| Specific membrane resistance ($k\Omega\text{cm}^2$) | 14.29 | 14.29 | 14.29 | 14.29 |
| Specific membrane capacitance ($\mu\text{F}/\text{cm}^2$) | 1 | 1 | 1 | 1 |
| Membrane time constant (ms) | 14.29 | 14.29 | 14.29 | 14.29 |
| Axial resistivity ($\Omega\text{-cm}$) | 200 | 200 | 200 | 200 |
| $I_{Na} g_{\max}$ ($\text{S}/\text{cm}^2$) | 2 | 0.72 | 2 | 0.85 |
| $I_{Kdr} g_{\max}$ ($\text{S}/\text{cm}^2$) | 0.03 | 0.008 | 0.1 | 0.01 |
| $I_d g_{\max}$ ($\text{S}/\text{cm}^2$) | 0.027 | 0 | 0.045 | 0 |
| $I_{\text{adap}}, g_{\text{adap}}$ (pS) | 800 | 0 | 10 | 0 |

##### Delayed rectifier potassium current

Delayed rectifier potassium currents (*I*_*Kdr*_) are known to contribute to action potential repolarization in numerous neuronal subtypes of the brain (Locke and Nerbonne, 1997; Murakoshi and Trimmer, 1999; Guan et al., 2007; Liu and Bean, 2014). We modeled this current in the LR neuron model using the channel gating properties of delayed rectifier potassium currents found in RS neurons (Liu and Bean, 2014). The channel model consists of 2 activation gates (Golomb et al., 2007) and no inactivation gates. The channel’s activation time constant (Liu and Bean, 2014) was tuned such that the model’s spike half width matches the experimentally obtained values. The channel was distributed in all 8 compartments of the model, and their *g*_*max*_ values are given in Table 2. The equations for voltage dependence of steady state activation (*n*_∞_) and the activation time constant (*τ*_*n*_) of the channel is described below.

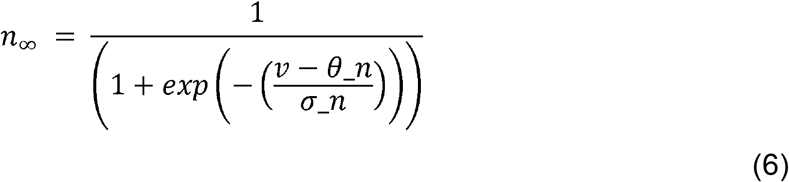

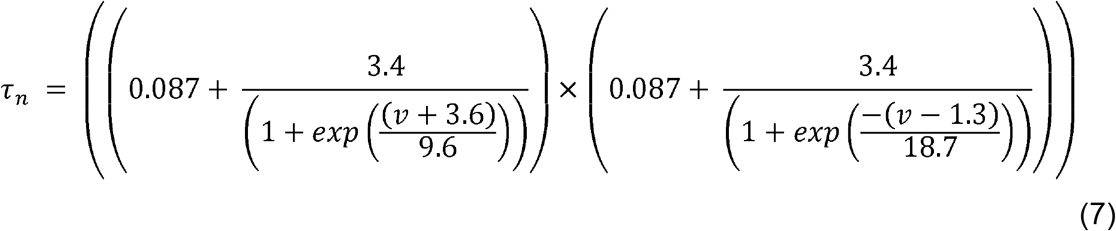

where *θ*_*n*_ = −20 *mV*, *σ*_*n*_ = 10.4 *mV*

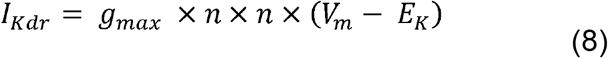

##### K_v_1 current

The K_v_1 current (also known as the d-current, (*I*_*d*_)) is a potassium current that is widely known to cause a delay to first action potential in many neuronal subtypes (Storm, 1988) (Goldberg et al., 2008; Kurotani et al., 2013). We modeled this current to capture the late spiking property of LR neurons that is observed in our physiological data. Similar to the fast sodium and delayed rectifier currents, this current was modeled using the Hodgkin Huxley formalism (Hodgkin and Huxley, 1952). Based on experimental data (Wu and Barish, 1992) and a previously published model (Golomb et al., 2007), the K_v_1 channel was modeled with 3 activation gates with faster kinetics and a slowly inactivating gate. This channel was distributed only in the somatic compartment of the neuron. The channel’s *g*_*max*_ is given in Table 2. The voltage dependence of steady state activation/inactivation (*a*_∞_, *b*_∞_) and their respective time constants (*τ*_*a*_, *τ*_*b*_) of the channel are given below.

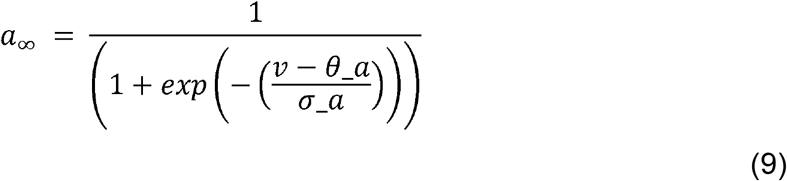

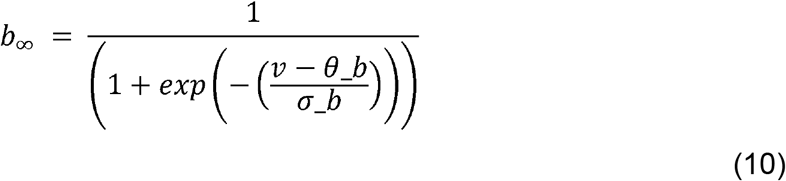

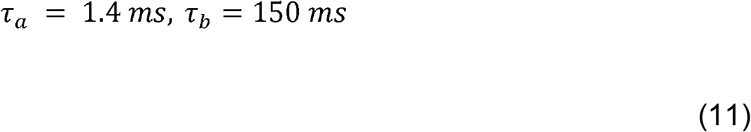

where *θ*_*a*_ = −50 *mV*, *σ*_*a*_ = 20 *mV*, *θ*_*b*_ = −70 *mV*, *σ*_*b*_ = −6 *mV*

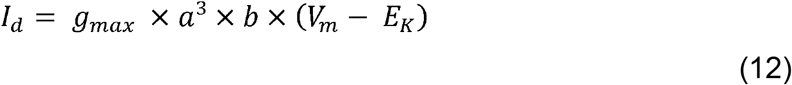

##### Adaptation current

Spike frequency adaptation was modeled using a linear mechanism as (*I*_*adap*_) described in previous studies (Treves, 1993; Fuhrmann et al., 2017). *I*_*adap*_ was modeled using the following equations (Treves, 1993).

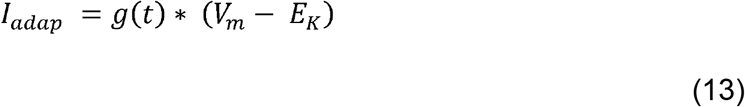

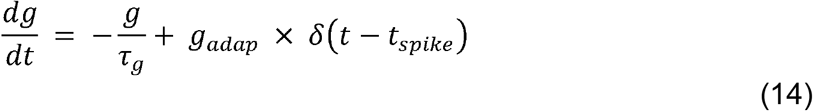

Briefly, when a cell fires an action potential, *g*(*t*) is increased by *g*_*adap*_, which decays to zero with a time constant of *τ*_*g*_.*t*_*spike*_ is the time at which the neuron spikes, and *E*_*K*_ is the potassium reversal potential. *g*_*adap*_ = 10 *pS* and *τ*_*g*_ = 500 *ms* (Liu and Wang, 2001).

### RS neuron model

#### Morphology and passive properties

The RS neuron model in our study consists of a somatic compartment and 21 dendritic compartments. The model’s input resistance is 150 MΩ and input capacitance is 95.3 pF. The model has a membrane time constant of 14.29 ms. The model’s resting membrane potential is −74.86 mV. Thus, the model’s passive properties accurately replicate those of RS neurons in layer 2/3 of RSG. The model’s ion channels and active properties are described in detail below.

#### Active properties

Similar to the LR neuron model, the RS neuron model has 3 voltage gated currents and a current for spike frequency adaptation (*I*_*adap*_). The voltage gated currents incorporated in the RS neuron model are fast sodium current (*I*_*Na*_), delayed rectifier potassium current (*I*_*Kdr*_) and K_v_1 current (*I*_*d*_). The model has a spike half width of 0.87 ms and a spike threshold of −54.9 mV. The spike frequency adaptation ratio of the model is 2.85, closely matching the experimental values.

##### Fast sodium current

The fast sodium current of the RS neuron model was modeled using Hodgkin Huxley’s equations (Hodgkin and Huxley, 1952). The channel’s voltage dependence of steady state activation/inactivation and their time constants were modeled using equations 1–5 (Martina and Jonas, 1997). The channel was distributed both in the somatic and dendritic compartments whose *g*_*max*_ values are described in Table 2.

##### Delayed rectifier potassium current

The delayed rectifier potassium current (*I*_*Kdr*_) was modeled based on the channel gating properties of K_v_2 currents found in RS neurons (Liu and Bean, 2014). Similar to the LR neuron model, the channel consists of 2 activation gates and does not exhibit any inactivation (Liu and Bean, 2014). The channel was placed in the somatic and dendritic compartments of the model (see Table 2 for *g*_*max*_ values). The channel equations are given below.

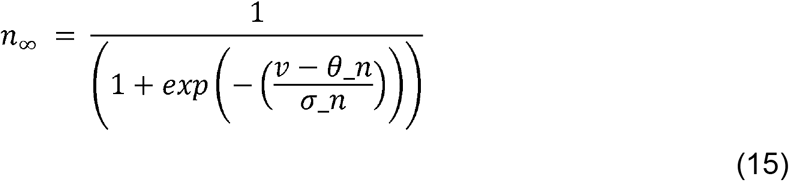

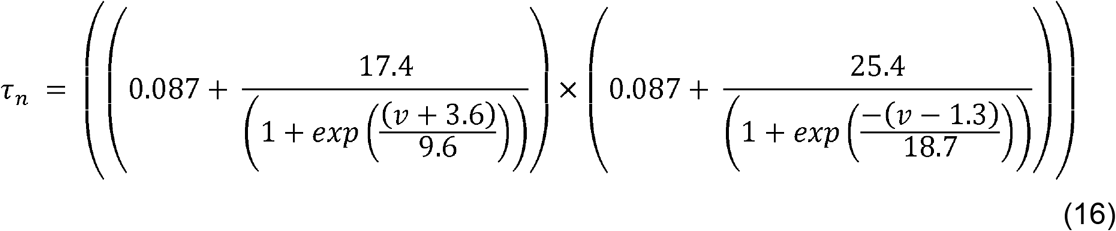

where *θ*_*n*_ = −20 *mV*, *σ*_*n*_ = 10.4 *mV*

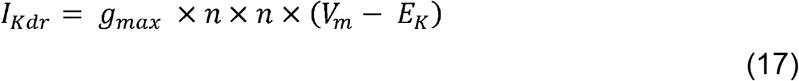

##### K_v_1 current

In order capture the observed late spiking behavior of layer 2/3 RS neurons of the RSC, *I*_*d*_ was also modeled in the RS neuron model. The channel’s gating mechanisms were modeled using equations 9–12 (Golomb et al., 2007). *I*_*d*_ was distributed only in the somatic compartment of the model (see Table 2 for Hcr *g*_*max*_ values).

##### Adaptation current

The spike frequency adaptation in the RS neuron model was modeled using the same schema (*I*_*adap*_) as described for LR neurons (equations 13–14) (Treves, 1993; Fuhrmann et al., 2017). The following parameters were used for this current: *g*_*adap*_ = 800 *pS* and *τ*_*g*_ = 500 *ms* In a subset of simulations (Figure 9) the adaptation current in RS cells was explicitly removed to study the contributions of adaptation to RS input-output transformations.

#### Synaptic inputs

The LR and RS neuron models received background and burst synaptic inputs. Briefly, the synaptic inputs were simulated in the following way: An AMPA and GABA synapse was placed in each of the dendritic compartments of both models. The location of the synapses in the respective branches varied across trials. Both the background and burst synaptic inputs were uniformly distributed over the dendritic tree of LR and RS neuron models in each trial. The properties of background and burst inputs are described in detail below.

#### Background inputs

The RS and LR neuron models received 10 AMPAergic background inputs. The time course of synaptic conductance of these background inputs is given by the following equation (Sterratt et al., 2011),

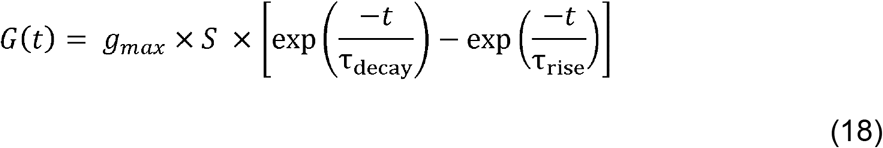

where *τ*_*decay*_ and *τ*_*rise*_ represent decay and rise time constant, respectively. *g*_*max*_ is the maximal synaptic conductance, and S is a normalization factor that equalizes the maximum of *G*(*t*) to *g*_*max*_. The values of *τ*_*rise*_ and *τ*_*decay*_ were 0.5 ms and 2.5 ms, respectively. The AMPAergic background inputs were modeled as Poisson spike trains with a frequency of 5 Hz and reversal potential of 0 mV (*E*_*AMPA*_).

Similarly, phasic GABAergic inputs were simulated for both models using equation 18. The *τ*_*rise*_ and *τ*_*decay*_ values for these inputs are 0.88 ms and 9.4 ms, respectively (Neymotin et al., 2011). Similar to excitatory background inputs, inhibitory inputs were simulated at a frequency of 5 Hz and reversal potential of −80 mV (*E*_*GABA*_).

The *g*_*max*_ values of the excitatory and inhibitory background inputs were chosen to capture the low background firing rates of pyramidal neurons observed *in vivo* and the ratio of excitatory-inhibitory (E-I) synaptic input strength (Xue et al., 2014) of neurons in the superficial layers of the cortex. For the LR neuron model, the *g*_*max*_ values of phasic excitatory and inhibitory background inputs were set to 2 nS and 12 nS, thereby maintaining an E-I ratio that is seen in experiments (Xue et al., 2014). Similarly, for the RS neuron model, the *g*_*max*_ values of phasic excitatory and inhibitory background inputs were set to 3 nS and 18 nS, respectively. The LR and RS neuron models have a background firing rate of ~1 Hz (Dégenètais et al., 2002; Koga et al., 2010; Nakamura et al., 2012).

#### Burst inputs

In addition to receiving background synaptic inputs, the models also received synchronous and identical burst inputs of various durations (25 ms, 50 ms, 100 ms, 200 ms, 500 ms, 2000 ms). The LR and RS neuron models received stimulation from 10 synchronous AMPAergic burst inputs. The time course of synaptic conductance of burst inputs were modeled using equation 18. The *τ*_*decay*_ and *τ*_*rise*_ of these inputs was set to 0.5 ms and 2.5 ms, respectively. The strength of burst inputs (*g*_*max*_) were set to 1200 pS for both models. For each burst condition (duration), the models were run for 300 trials.

#### Data analysis of model results

The models were simulated using NEURON 7.5 simulation environment (M.L Hines and N.T.Carnevale, 2001) with an integration time step of 0.025 ms. Simulation output was written into binary files and analyzed using custom programs written in MATLAB (R2018b) software. Spike threshold, spike half width, input resistance, membrane time constant and input capacitance of the models were calculated using the same method that was used for experimental data.

Signal to noise ratio (SNR) in response to the burst input was calculated using the following formula (Duguid et al., 2012),

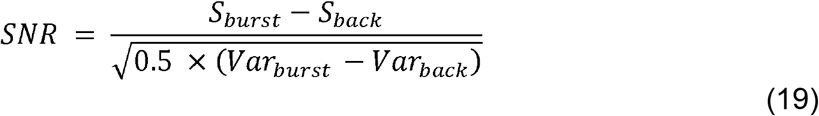

*S*_*burst*_ is the average number of spikes in ‘x’ ms post burst onset, where x = burst duration+125 ms. *S*_*back*_ is the average number spikes per ‘x’ ms from 4000 ms to 8000 ms post burst onset. *Var*_*burst*_ and *Var*_*back*_ are the corresponding variances of the number of spikes during those two time intervals.

The spike timing precision (jitter) in response to the burst input was computed by calculating the median absolute deviation of spike latencies from all trials. Bootstrapped resampling was then used to compute a distribution of jitter values, and significance between the jitter of LR and RS neurons was established by 2-tailed t-test (95% confidence interval) (Sudhakar et al., 2015). Significance in the probability of spiking between LR and RS neurons was established by Pearson chi-square test (Alan Agresti, 2007) (95% confidence interval).

#### In vivo dataset related modeling

In order to determine if LR and RS neuron models can sustain continuous firing as would be expected from the firing of head direction neurons in the preferred direction during motionless conditions, we stimulated the LR and RS neuron models with spike trains of neurons recorded from the post subiculum that had one preferred head-direction angle (head-direction cells) of awake mice (Peyrache et al., 2015). Spike data was downloaded from the website of CRCNS (https://crcns.org/data-sets/thalamus/th-1/about-th-1; Peyrache, A., Buzsáki, G. (2015) Extracellular recordings from multi-site silicon probes in the anterior thalamus and subicular formation of freely moving mice. CRCNS.org. http://dx.doi.org/10.6080/K0G15XS1) and given as input to the neuron models. Briefly, the LR and RS neuron models were stimulated with 10 synchronous input spike trains of head direction neurons recorded from post subiculum. Simulations were run for one entire awake epoch in the th-1 dataset, 1200 seconds in duration (Peyrache, A., Buzsáki, G. (2015) Extracellular recordings from multi-site silicon probes in the anterior thalamus and subicular formation of freely moving mice. CRCNS.org. http://dx.doi.org/10.6080/K0G15XS1). The simulations were repeated for 30 trials each. SNR was calculated according to equation 19. The resulting SNR was binned and plotted as a function of stimulus duration.

## ACKNOWLEDGEMENTS

We would like to thank Wayne Aldridge, Barry Connors, Scott Cruikshank and Vaughn Hetrick for their comments on the manuscript and Vaughn Hetrick for his technical assistance. This work was supported by an NSF graduate research fellowship to EKWB and by grants to OJA from the NIH (R03MH111316), the American Epilepsy Society Junior Investigator Award, the Massey Foundation and the MICDE Catalyst Award.

